# CRISPR/Cas9 targeting Ttc30a mimics ciliary chondrodysplasia with polycystic kidney disease

**DOI:** 10.1101/2020.11.27.400994

**Authors:** Maike Getwan, Anselm Hoppmann, Pascal Schlosser, Kelli Grand, Weiting Song, Rebecca Diehl, Sophie Schroda, Florian Heeg, Konstantin Deutsch, Friedhelm Hildebrandt, Ekkehart Lausch, Anna Köttgen, Soeren S. Lienkamp

## Abstract

Skeletal ciliopathies (e.g. Jeune syndrome, short rib polydactyly syndrome, Sensenbrenner syndrome) are frequently associated with cystic kidney disease and other organ manifestations, but a common molecular mechanism has remained elusive.

We established two models for skeletal ciliopathies (*ift80* and *ift172*) in *Xenopus tropicalis*, which exhibited severe limb deformities, polydactyly, cystic kidneys, and ciliogenesis defects, closely matching the phenotype of affected patients.

Employing data-mining and an *in silico* screen we identified candidate genes with similar molecular properties to genetically validated skeletal ciliopathy genes. Among four genes experimentally validated, CRISPR/Cas9 targeting of *ttc30a* replicated all aspects of the phenotypes observed in the models of genetically confirmed disease genes, including ciliary defects, limb deformations and cystic kidney disease.

Our findings establish three new models for skeletal ciliopathies (*ift80*, *ift172*, *ttc30a*) and identify TTC30A/B as an essential node in the network of ciliary chondrodysplasia and nephronophthisis-like disease proteins implicating post-translational tubulin modifications in its pathogenesis.

## Introduction

Primary cilia are hair-like appendages that extend from the surface of most differentiated cells. Their functional impairment or loss results in ciliopathies^1,2^. Ciliopathies can affect a range of organ systems and tissues, including the kidneys, liver, eyes, testes, brain and skeleton. Often, multiple organ manifestations occur in combination (e.g. Bardet-Biedl Syndrome affecting kidney, brain, eyes, gonads and digit number), but depending on the affected locus, only certain combinations of phenotypes may co-occur^3,4^.

Most genes affected by mutations in ciliopathy patients encode for structural components of the primary cilium^1–3^. Some protein groups co-localize to ciliary subcompartments, and are associated with common clinical features. Examples are the Bardet-Biedl related BBSome components at the base of cilia^5^, proteins in the proximal cilium that are mainly linked to nephronophthisis^6^, or components of the axonemal intraflagellar transport (IFT) machinery linked to skeletal ciliopathies (SC) such as Jeune’s asphyxiating thoracic dysplasia (JATD), Mainzer-Saldino syndrome (MZSDS) and Sensenbrenner syndrome or Cranioectodermal dysplasia (CED)^4,7,8^. However, many ciliopathy proteins don’t adhere to these categories. For example, mutations of basal body genes (*CEP120*) were identified in SC patients^4,9^, and IFT components (*IFT27, IFT74*) can cause Bardet Biedl syndrome^10,11^. Thus, the molecular cause of the variability and clinical overlap of certain ciliopathy manifestations is elusive.

Despite recent advances in genetics that identified most genes affected in ciliopathies, and studies of the ciliary proteome in cell culture models, the understanding of ciliary protein compositions and their function in each specific tissue remains incomplete^12–14^. Posttranslational modifications (PTMs) of axonemal tubulin by polyglycylation and - glutamylation contribute to the “tubulin code” that diversifies tubulin function and supports the structural integrity of cilia^15–17^. *In vitro* studies have shown that ciliary polyglutamylation by TTLL5 and TTLL6 depends on the Joubert syndrome associated proteins ARL13B and CEP41^18,19^, but if tubulin PTMs are defective in other ciliopathies or contribute to the phenotypic pleiotropy remains unknown.

One subgroup of ciliopathies affects patients with skeletal deformities^4,9^. Typical phenotypes of SCs are shortened ribs resulting in a narrow thorax, truncated limbs (micromelia), shortened fingers (brachydactyly), and multiple digits (polydactyly). In some cases, the kidneys (nephronophthisis or cystic kidneys), eyes (retinopathy), liver function (liver fibrosis, liver cysts), or left-right axis formation (*situs inversus*) can be affected.

The Hedgehog (Hh) signaling pathway is one of many cellular signals that is frequently found to be disrupted by defective ciliary function^20–32^. The ciliary Hh pathway is required for chondrocyte proliferation, differentiation, and limb patterning^33,34,35,36^. Two of the best characterized SC genes are *ift80*^37–39^ (mutated in SRPS III and IV and JATD) and *ift172*^40–42^ (mutated in JATD and MZSDS). Both are IFT-B subcomplex proteins^43^ and although genetic interactions were described, they do not interact physically with each other^40,43,44^. In model organisms, biallelic loss-of-function mutations result in shortened, missing or fewer cilia. Morphant and mutant zebrafish larvae have pronephric glomerular cysts, retinal degeneration, pericardial edema, slightly smaller eyes, and a curved tail^39,40,45^. In addition, *ift172* morphants have otolith defects, hydrocephalus and craniofacial cartilage defects^40^. In mice, null alleles of *ift80* or *ift172* are prenatally lethal^20,24^, while mice with hypomorphic alleles have a low survival rate and suffer from shortening of long bones, a narrow thorax and polydactyly^24,46,47^. Chondrocyte specific loss-of-function of either gene in conditional knockout mice results in severely shortened limbs^48,49^. Conditional Col2α1; IFT80^fl/fl^ mice revealed that chondrocytes are disorganized, the growth plate is smaller, and articular cartilage is thickened^48,50^. Loss of *ift80* or *ift172* reduces the expression of Hedgehog target genes, such as *patched1* or *gli1*^24,47^, in line with the hypothesis that Ift80 acts downstream of Smoothened as its agonist SAG and the second messenger Gli2 can rescue the loss of *Ift80*^48^. Embryonic renal dysplasia has been observed in mice with hypomorphic alleles of *ift172*, but no studies have addressed their role in kidney function.

Mutations in at least 27 known disease genes currently account for most genetic diagnoses in cases of SC^4^. However, because SC is a rare genetic disorder, the known causal genes might not cover the complete spectrum of genes relevant to the pathophysiology of SC. A more complete understanding of the molecular network of SC proteins is needed to shed light on its pathogenesis and offer potential therapeutic options.

Here we describe two novel models for chondrodysplasia using CRISPR/Cas9 mediated editing of *ift80* and *ift172* in *Xenopus*. CRISPR targeted froglets developed severe limb defects during metamorphosis with shortened limbs due to chondrocyte differentiation defects, syn- and polydactyly, and polycystic kidneys, phenocopying major clinical features of ciliary chondrodysplasia. Impaired ciliogenesis was detected in multiciliated cells and resulted in defective fluid excretion during tadpole stages. Using a data-mining approach, we identified candidate genes with similar molecular properties to the established disease genes. Among these, only *ttc30a* targeting mimicked the full spectrum of chondrodysplasia phenotypes, i.e. impaired ciliogenesis, fluid retention in tadpoles, polycystic kidneys, and strongly affected limb development, including polydactyly in froglets. Thus, Ttc30a acts in ciliary signaling disrupted in chondrodysplasia and cystic kidney disease. Enhanced expression in chondrocytes and osteocytes of Ttc30a1 and Ttc30b suggest an evolutionarily conserved role in mammals. Together, our findings identify TTC30A/B as a critical node in the pathogenesis network of ciliary chondrodysplasia with polycystic kidney disease.

## Results

### CRISPR/Cas9 targeting of *ift80* and *ift172* leads to chondrodysplasia and cystic kidneys in *Xenopus tropicalis*

We chose *Xenopus* to establish a model for ciliary chondrodysplasia, because it is an aquatic organism that allows for analysis of limb development, has a high sequence conservation to humans, is an established model in cilia research, and provides good experimental access^50–54^. We targeted two well characterized SC genes (*ift80* and *ift172*) for loss-of-function analysis using CRISPR/Cas9 based genome editing. Indel frequency and knockout scores were used to identify highly effective sgRNAs against both targets in *X. tropicalis* embryos (Supplement Fig. 1 a-c). Injection of F0 embryos with Cas9/sgRNA ribonucleoprotein (RNP) resulted in mosaic gene disruption, and has previously been shown to elicit specific phenotypes at high frequency and penetrance^55–60^.

Phenotypic analysis of 3-4 day old embryos (stage 45) injected at the one-cell stage showed prominent generalized edema in both *ift80* and *ift172* targeted embryos. Importantly, this phenotype was rescued by co-injecting *ift80* or *ift172* mRNA, demonstrating the specificity of the sgRNA mediated loss-of-function (Fig. 1 a, b). Morpholino oligonucleotide knockdown in *X. tropicalis* and CRISPR/Cas9 targeting elicited the same phenotype in *X. laevis* confirming the specificity of the phenotype in a separate species (Supplementary Fig. 1 d-j).

**Figure 1:**
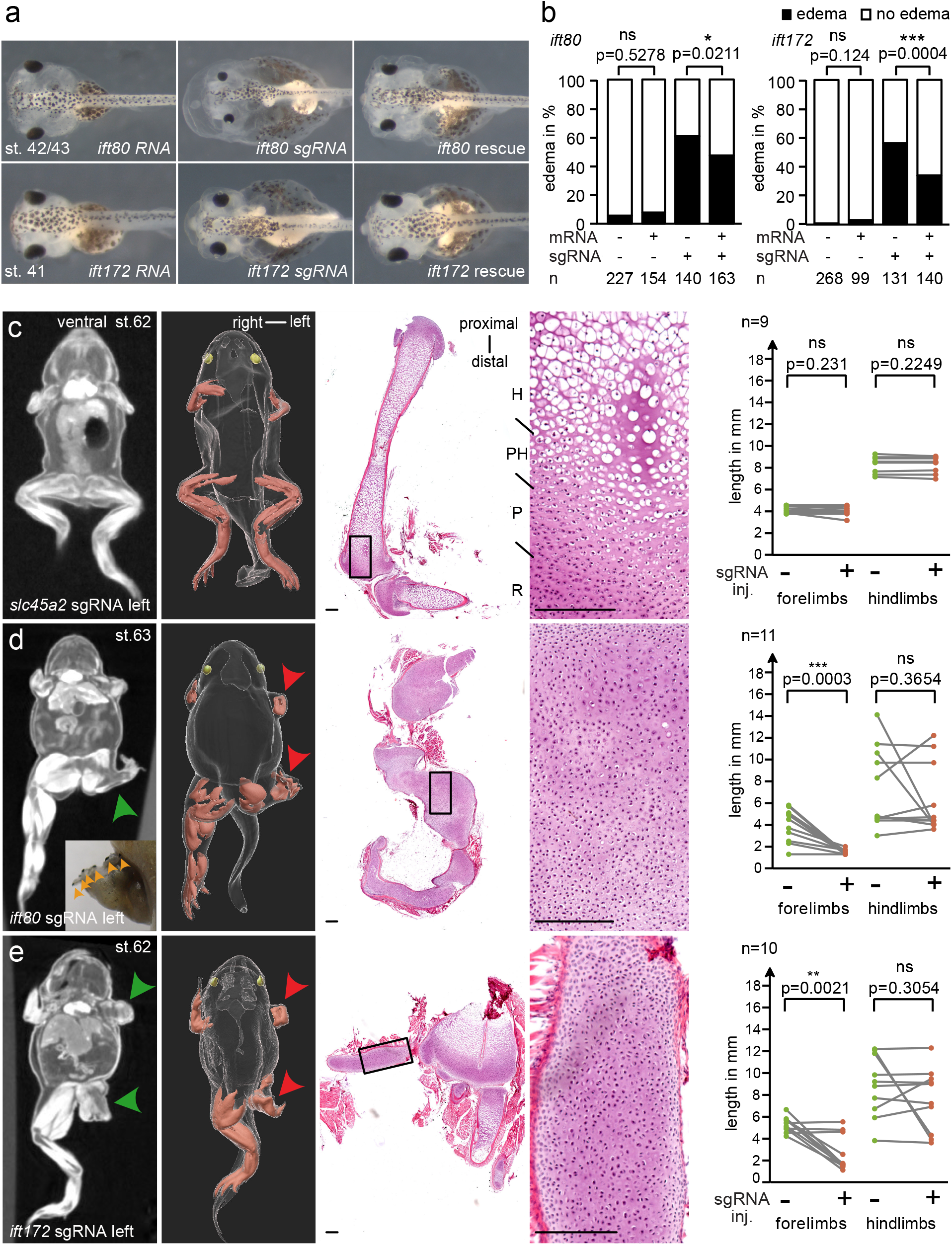
CRISPR targeting of *ift80* and *ift172* leads to skeletal defects in *X. tropicalis*. (a, b) Analysis of edema formation in *ift80* and *ift172* mutants at stage 41/42. Note the smaller eyes in *ift80* targeted embryos. (c - e) Limb phenotypes in post-metamorphic froglets. MicroCT scans of representative mutant froglets of *slc45a2*, *ift80* and *ift172* are shown in dorsal views. Injections of RNPs were performed unilaterally (left) at the 2-cell stage. For the same animals 3D-reconstructions are pictured. Limb muscles are shown in red, eyes in yellow. Histological sections of hindlimbs stained with H&E. Scale bars represent 200μm (c) *slc45a2* animals showed regular limb development. (d) Polydactyly is indicated by orange arrowheads to mark black nailed digits. MicroCT scans and histological stainings revealed cartilage accumulations in CRISPR-targeted froglets (green arrowheads). 3D-reconstructions demonstrate that only left limbs were affected (red arrowheads). Length of respective uninjected (green dots) vs injected (red dots) limbs of the same individual are plotted (grey lines). R - resting zone; P - proliferating zone; PH - pre hypertrophic zone; H - hypertrophic zone. p>0.05 ns (not significant); p<0.05 *; p<0.01 **; p<0.001 ***

Embryos targeted for *ift80* or *ift172* disruption sometimes had smaller heads and narrower eye width which often occurred in combination with head edema (Fig. 1 a; Supplementary Fig. 1d, e). Craniofacial defects also appeared at high rates in *X. laevis* CRISPR/Cas9 targeted embryos (Supplementary Fig. 1 g, h).

The most prominent phenotypes in chondrodysplasia patients are malformations of the axial skeleton and limbs, i.e. shortened ribs, long bone dysplasia, and polydactyly^4^. Unilateral CRISPR-targeting of F0 animals has been shown to be effective to disrupt genes essential for post-metamorphic limb development in *Xenopus*^61–63^. To test if Ift80 and Ift172 act in limb development in *Xenopus* we unilaterally injected two-cell stage embryos, which limits gene editing to only one half of the body while retaining the unaffected half as an internal control. We targeted *slc45a2* as an additional negative control, which results in a loss of pigmentation that can be readily observed at earliest stages and controls for the correct delivery and function of the Cas9/RNP-endonuclease^59,64^. Apart from the expected loss of skin pigmentation, metamorphic froglets were phenotypically normal, demonstrating that sgRNA/Cas9 injection *per se* did not have an impact on development (Fig. 1 c, Supplementary Fig. 4 j).

In contrast to controls, froglets that were CRISPR-edited at the *ift80* or *ift172* locus showed severe limb defects. Both fore- and hindlimbs were substantially shorter than in uninjected control or *slc45a2* targeted froglets (Fig 1 c, d, e, green and red arrowheads). In addition, limb diameters were clearly thickened compared to controls.

In most animals the fingers and toes were shortened and less developed, sometimes not even identifiable (Fig. 1 d, e). The number and size of clawed digits of the hindlimbs varied substantially from wildtypes, in particular in *ift80* targeted froglets. *X. tropicalis* usually possess six digits at the hindlimb, four of which form claws^65^. In *ift80* targeted froglets, the number of clawed toes were consistently either reduced to less than three (syndactyly) or increased to six or seven (polydactyly) (Fig. 1 d). In conclusion, this phenotype is highly reminiscent of the polydactyly phenotype observed in mouse models and human patients^24,37,40,49,66^.

In order to explore the phenotypic changes in more detail, we conducted microCT scans of mosaic mutant froglets. There was no difference between injected and uninjected sides of the control *slc45a2* targeted froglets, the length measurements were much less variable than in *ift80* and *ift172* (Fig. 1 c-e). Despite unilateral RNP injections, we observed bilateral limb malformation in some animals, suggesting that in these cases, the first cell cleavage had not fully separated at the time of injection, allowing for gene editing to occur across the midline. Comparing limbs of *ift80* and *ift172* disrupted froglets on the injected versus the uninjected side, we detected a strong shortening of forelimbs and a less pronounced length difference of the hindlimbs (Fig. 1 c-e). Iodine based contrast enhancement allowed us to distinguish muscle, cartilage, and other internal organs based on density differences. We found that the shortened, not yet ossified limbs consisted mainly of cartilage accumulations (Fig 1 d, e, green arrowheads), consistent with the phenotype in humans and mouse models^48,50,67^. Histological sections confirmed that the resting and proliferation zones of cartilage dominated clearly in *ift80* and *ift172* targeted animals at the expense of the hypertrophic and ossification zones visible in *slc45a2* targeted control animals (Fig. 1 c-e). In general, the chondrocytes were less well organized in *ift80* and *ift172* depleted cartilage than in limbs of controls. Similar phenotypes have been described for conditional *ift80* knockout mice and other chondrodysplasia models^24,26,68–70^.

Contrast enhanced microCT was also able to resolve the structure of the mesonephric kidneys in the post-metamorphic froglets at high resolution and revealed polycystic kidneys in most of the mutant animals (Fig. 2 a-c). Quantification and volumetric analysis showed both *ift80* and *ift172* disruption resulted in a significantly higher number of cysts (> 0.2mm) in comparison to slc45a2 controls, particularly on the injected side (Fig. 2 d). The occurrence of cysts on the uninjected side in *ift80* mosaic mutants likely reflects the incomplete segregation of gene editing to one half, as observed for limb measurements. The total cyst volume was larger on the injected side of *ift80* but not significantly changed for *ift172* edited froglets (Fig. 2 e). The volume of non-cystic renal parenchyma was not significantly different (Fig. 2 f). To our knowledge, this is the first report of polycystic mesonephric kidneys in *Xenopus*.

**Figure 2:**
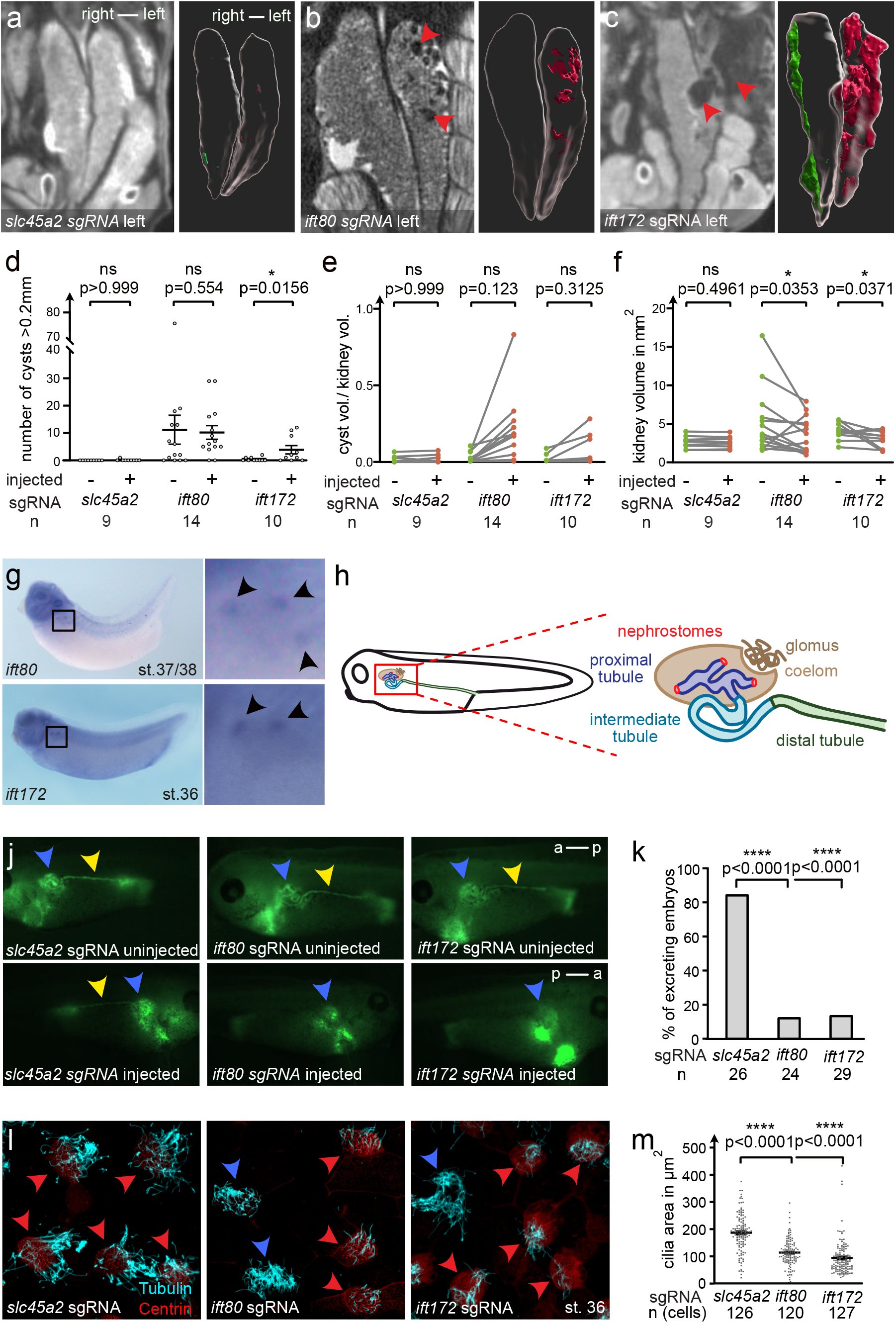
Cystic kidney disease and cilia defects in *ift80*- and *ift172* CRISPR targeted *X. tropicalis*. (a - c) CRISPR/Cas9 targeting of *slc45a2*, *ift80* and *ift172* were performed unilaterally in two-cell-stage embryos. Mesonephri of st. 61-63 froglets were analyzed by microCT scans and kidneys and cysts (red arrowheads in b, c) were segmented for 3D volumetric analysis (a – c; red - injected; green - uninjected). (d) The number of cysts (> 0.2mm), (e) the ratio of total cyst volume (of cysts > 0.2mm) to kidney volume and (f) the total kidney volume (excluding cysts > 0.2mm) was calculated and compared between right untreated (−) and left injected (+) kidneys. (g) Whole mount *in situ* hybridization detects *ift80* and *ift172* in the multiciliated nephrostomes of the pronephros in *X. laevis*. (h) Schematic depiction of the embryonic renal system of *Xenopus*. (j, k) Excretion assay with fluorescein-dextran at stage 38-40. Blue arrowheads point to the proximal part of the pronephros. Yellow arrowheads indicate fluorescence signal in the distal tubule, lacking on the injected side (j). a – p: anterior – posterior. (l) Confocal images of multiciliated epidermal cells (MCCs) stained against acetylated tubulin (cyan). Centrin-RFP fusion protein served as a lineage marker (red arrowheads) and indicates CRISPR-targeted MCCs. Blue arrowheads point to non-targeted (wild-type) cells. (m) The ciliated area was determined for each cell. Error bars indicate SEM. p>0.05 ns (not significant); p<0.05 *; p<0.01 **; p<0.001 ***

We conclude that froglets with mutations in *ift80* and *ift172* displayed shortened extremities with polydactyly and cystic kidney disease, confirming that chondrodysplasia can be modelled in *Xenopus*. This supplements the available mouse and zebrafish lines, and offers new opportunities to analyze chondrodysplasia genes and their prospective functions^49,50,67,71^.

### *Ift80* and *ift172* targeting affects multiciliated cells resulting in pronephric excretion defects

Next, we explored the effect of *ift80* and *ift172* disruption on a cellular level. The most prominent phenotype of *ift80* and *ift172* mutant tadpoles was generalized edema. Suggesting a deficiency in water or ion homeostasis due to pronephric, lymphatic, vascular, or cardiac insufficiency. *In situ* hybridization of established marker genes (*aplnr*, vasculature; *nkx2.5*, heart; *prox1*, lymphatic system) did not reveal an obvious developmental disruption of any of these organ systems in *Xenopus* embryos targeted for *ift80* and *ift172* disruption (Supplementary Fig. 2 c, d). Given the prominent renal phenotype in froglets, the edema phenotype in embryos, and because Jeune syndrome patients and mouse models frequently suffer from kidney disease, we focused on the pronephros of *X. tropicalis* embryos47. We did not find structural tubular defects when staining for the tubular epithelium (Tomato lectin; Supplementary Fig. 2 e, f) nor were the expression of segment-specific marker genes (*nkcc2*, distal tubule; *sglt1* proximal tubule) affected (Supplementary Fig. 2 c, d) suggesting a normal differentiation of tubular tissue.

Expression analysis of *ift80* and *ift172* by *in situ* hybridization revealed both genes to be prominently expressed in the nephrostomes (Figure 2 g). These represent the most proximal segments of the pronephric tubule that contain densely ciliated cells which facilitate uptake of coelomic fluid into the pronephric tubule (Figure 2 h). To test whether impaired fluid uptake into the tubule could explain the observed edema phenotype in *ift80* and *ift172* bilaterally disrupted embryos, we employed a dextran excretion assay. Embryos were injected with sgRNA/Cas9 at the two-to four-cell stage, and fluorescently labeled dextran was injected into the coelom at stage 39-40 when the pronephros is fully functional. Time-lapse movies of untreated or *slc45a2* sgRNA injected (negative control) embryos showed rapid uptake and acceleration of dextran through the pronephric tubule and excretion at the cloaca on both sides in 84,6% of embryos (Fig. 2 j, k). In contrast, entry into the pronephric tubule was observed in only 12,5% and 13,8% of embryos within 3 min on the *ift80* and *ift172* targeted side (Fig. 2 j, k), respectively. Dextran uptake was comparable to wildtype and much faster on the non-targeted side of unilaterally gene edited embryos. Normal excretion on the uninjected side could also be indirectly observed in the time-lapse movies where dextran entered the medium of *ift80* and *ift172* targeted embryos (Supplementary Movie 1).

Motile cilia in the nephrostomes actively propel fluid into the renal tubules. Thus, structural defects of ciliated cells, which has been described for *ift80* and *ift172* mutant zebrafish, may explain the observed loss of excretion in CRISPR targeted tadpoles^45,71^. To analyze the role of *ift80* and *ift172* in motile cilia, we turned to the epidermal multiciliated cells, a well established and accessible model for ciliogenesis. *Centrin-RFP* mRNA was used as an injection marker to identify ciliated cells that received the CRISPR RNPs and label basal bodies of ciliated cells. The mosaic distribution allowed us to analyze CRISPR targeted and wildtype cells in the same images. Cilia appeared to be less and shorter in length which was confirmed by quantification of cilia in *ift80* and *ift172* knockout cells. No such difference was observed in *slc45a2*-targeted controls (Fig. 2 l, m). In contrast, basal body number and apical docking was not altered in crispant MCCs (multiciliated cells) when compared to controls (Supplement Fig. 2 g). Thus, the ciliogenesis defect is consistent with findings in other model organisms^45,71^ and a likely cause for impaired tubular uptake.

To gain more detailed insights into the molecular signals affected by a loss of *ift80* or *ift172*, we investigated the transcriptional profile of embryos targeted for *ift80* and *ift172*. Because embryos were *in vitro* fertilized and developed synchronously, stage dependent variability was minimized between control and CRISPR targeted embryos. We isolated DNA and confirmed a high indel frequency in the same samples subjected to RNA-Seq analysis at stage 22 (neurula) (Suppl. Figure 3 a, b). The top most downregulated genes were *ift80* and *ift172*, respectively (Fig. 3 a, Suppl. Fig. 3 c, d, and Suppl. Tables 1 and 2), confirming the specificity of our targeting strategy and a direct effect on mRNA levels likely due to nonsense-mediated decay. In each condition, we found 15 (*ift80*) and 5 (*172*) genes to be differentially regulated (Suppl. Fig. 3 c, d). We found three genes (*abcb1* and two uncharacterized transcripts) to be significantly upregulated in both conditions (Fig. 3 a). The ATP-dependent multi-drug resistance transporter Abcb1 is transcriptionally regulated by Gli1 and β-catenin^72,73^. Interestingly, additional five genes that only reached significance in one condition were also associated with Hedgehog-signaling (*ccng1*, *ulk1*, *riok3*, *olfm4*, *col1a1*) and four with Wnt-signaling (*c3ar1*, *col1a1*, *olfm4* and *ulk1*) (Figure 3 a)^72,74–82^. Thus, we detect Hedgehog- and Wnt-dependent gene regulation to be altered upon depletion of *ift80* and *ift172* in *Xenopus* embryos.

**Figure 3:**
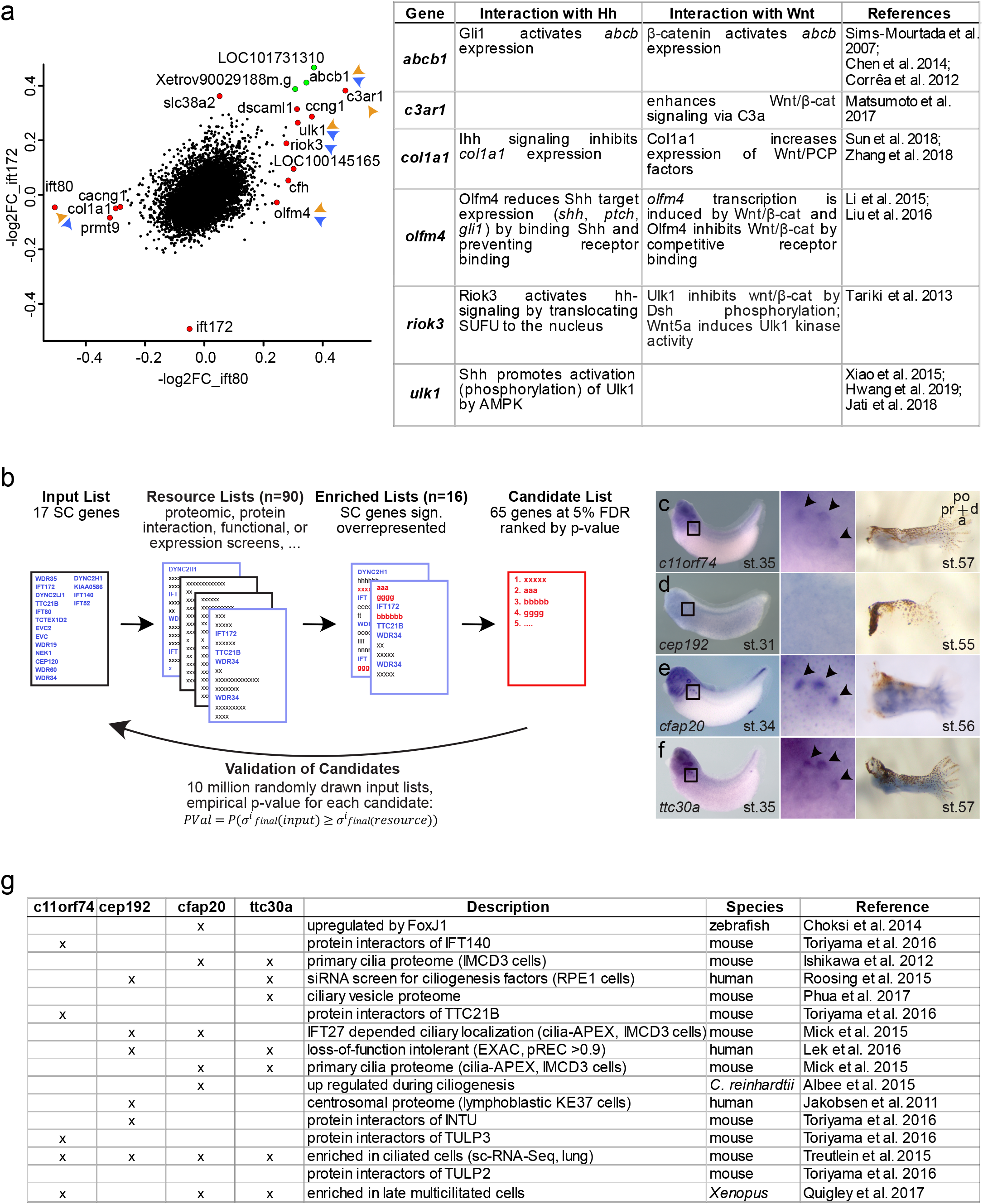
Transcriptomic profiling of *ift80* and *ift172* CRISPR targeted embryos and *in silico* screening for candidates with similar properties to identified SC-genes. (a) Scatterplot of the log2-fold changes in mRNA expression between *ift80*- and *ift172* targeted embryos. Significantly changed transcripts are labeled with gene names. Red dots mark genes exclusively altered in *ift80*- or *ift172* targeted embryos. Green dots indicate genes significantly changed in both conditions. Blue arrowheads point to genes for which an association with the hedgehog signaling pathway has been described, orange arrowheads on genes with wnt-associations. A table of respective associations and references thereof. (b) Schematic of the *in silico* screen. Known SC-disease genes serve as an input list. Resource lists represent datasets from various published screening approaches and are tested for enrichment of genes contained in the input list genes. Genes in significantly enriched resource lists are then scored based on membership or rank. The gene scores are statistically validated by an empirical p-value based on 10 million random drawings of input list. (c - f) Whole mount *in situ* hybridizations for four candidate genes (*c11orf74*, *cep192*, *cfap20*, *ttc30a*) that were experimentally followed up are shown for tadpoles (nephrostome expression enlarged) and limb buds. (g) Resource Lists in which the candidate genes occurred.

mRNA expression analysis in wildtype embryos of differentially regulated genes identified by RNA-Seq found an overlapping expression with chondrodysplasia genes in neuronal tissue and neural crest. This was particularly true for *ccng1*, *ulk1* and *riok3* (Supplement Fig. 3 e). However, we did not find a change in the Sonic Hedgehog marker *nkx2.2* by *in situ* after unilateral *ift80* or *ift172* targeting, suggesting that the differences in mRNA abundance are below the detection level of *in situ* hybridization (Suppl. Fig. 3 f, g).

### Data mining identifies skeletal ciliopathy associated genes

Next, we aimed to identify novel proteins related to the pathogenesis of SC. We reasoned that proteins with similar molecular properties, such as subcellular localization, common protein interactors, or differential regulation in loss-of-function models, could be unrecognized components of the pathogenetic molecular mechanism. Therefore, we assembled lists of genes from 90 published genome-wide screens or unbiased analyses (“resource lists”) (Suppl. Table 3) based on literature curation. Because these lists were obtained from screening experiments generated across various species, all gene identifiers were matched to the closest human ortholog. We identified 16 resource lists to be significantly enriched for genetically validated chondrodysplasia genes in comparison to 10 million random draws of an equal number of query genes (Figure 3 b). Within these 16 lists, 65 unique genes not yet linked to chondrodysplasia were found to be significantly enriched (Suppl. Table 4). Among these potential candidate genes were three that have been identified as *bona fide* chondrodysplasia genes in the meantime (i.e. *ift122*, *ift43* and *traf3ip1*^70,83,84^). *Ift22* was described to be mutated in a short rib-polydactyly syndrome (SRPS) type IV patient, *ift43* and *traf3ip1* in SRPS type II patients and *traf3ip1* additionally in a case of JATD^70,83,84^. This confirmed that our approach was able to predict valid genetic associations.

Given that the system worked well with chondrodysplasia genes, we used all gene sets annotated in phenotypic series in OMIM as input. A significant number of enriched lists could be identified for 13 disease entities, many of which were ciliopathies (e.g. Meckel syndrome, nephronophthisis, primary ciliary dyskinesia; Suppl. Table 6).

Among the 65 potential chondrodysplasia related candidate genes, we focused on genes that have not been linked to any ciliopathies nor characterized in a knockout mouse model. We focused on the top four genes with the lowest p-value, that were targetable in *X. tropicalis* (*C11ORF74*, *CEP192, CFAP20* and *TTC30B*) for functional analysis (Fig. 3 c-g). All proteins encoded by these four candidate genes had previously been associated with cilia. CFAP20 (cilia and flagella associated protein 20) regulates cilia length and morphology^85,86^. Entering of the BBSome into the cilium via interaction with the IFT-A complex is controlled by C11ORF74^87^. CEP192 (centrosomal protein of 192kDa) is required for mitotic centrosome and spindle assembly^88,89^. TTC30B (tetratricopeptide repeat domain 30b, also known as IFT70) is essential for polyglutamylation and polyglycylation of axonemal tubulin and is part of the IFT-B complex^90–92^. TTC30B has one closely related homolog in humans (*TTC30A*) and three in mice (*Ttc30a1*, *Ttc30a2*, *Ttc30b*), but only one ortholog in *X. tropicalis* and *Danio rerio* (*ttc30a/fleer)* (Fig. 4 a)^91^.

**Figure 4:**
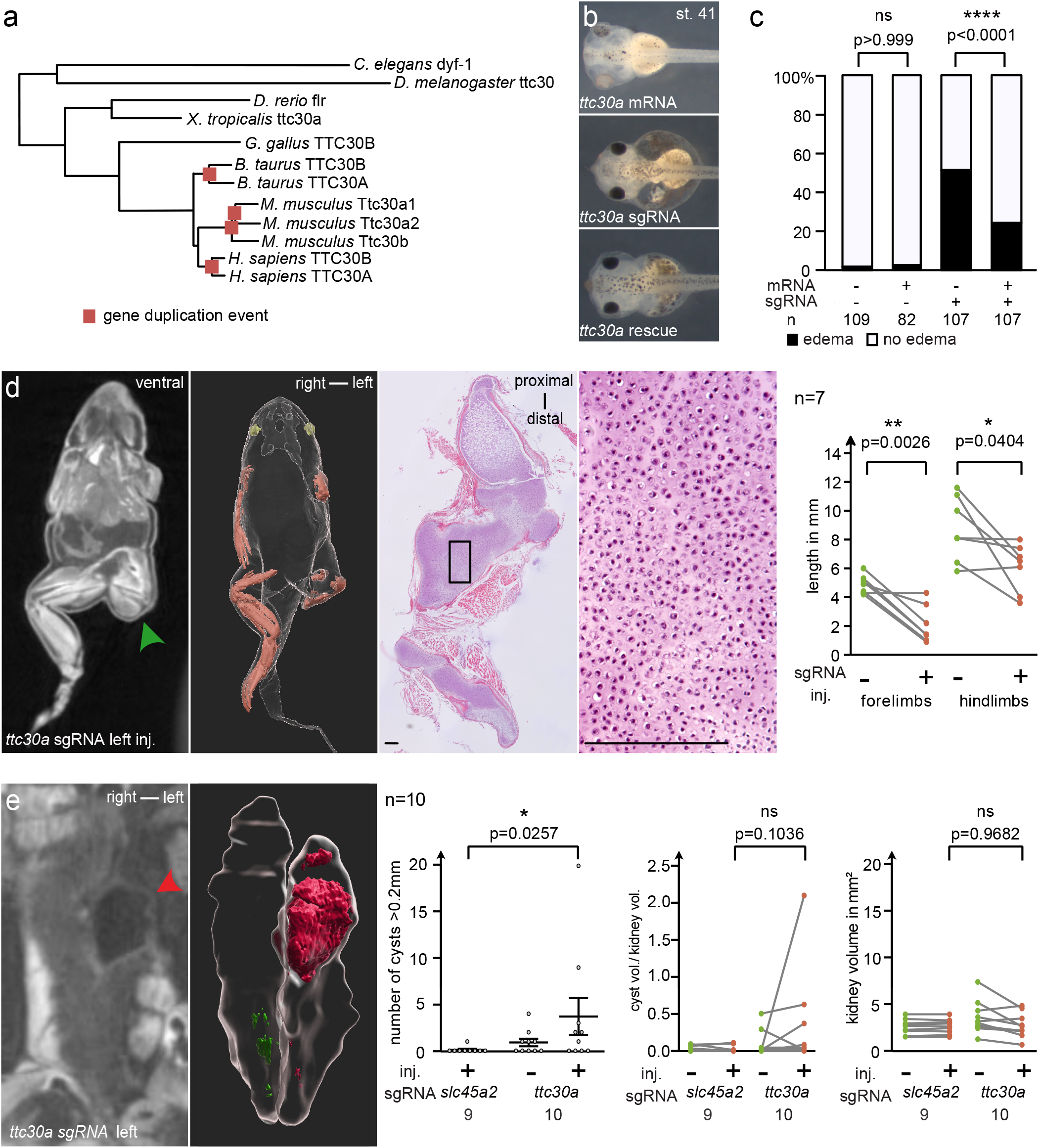
*ttc30* loss of function resembles the SC-phenotype. (a) Phylogenetic tree of protein sequences of TTC30A/B orthologs in various species. Genetic duplication events are marked by red squares. (b, c) CRISPR targeting of *ttc30a* at the one-cell stage led to edema formation, quantified in (c). Co-injection of *ttc30a* mRNA partially rescued the phenotype. (d) Unilaterally CRISPR targeted *ttc30a* froglets developed shortened limbs on the injected side (green arrowhead). MicroCT analysis and histological sections stained with H&E demonstrated accumulation of cartilage (green arrowhead; lines represent 200μm). 3D-reconstructions are of the same individuals. Quantification of limb length reduction of *ttc30a* targeted compared to non-targeted side. (e) MicroCT scans show cystic kidneys in *ttc30a* targeted animals (red arrowhead). 3D-reconstruction of the kidneys and cysts (red) were used for volumetric analysis. Quantifications of cyst number (> 0.2mm), the ratio of total cyst volume (of cysts > 0.2mm) to kidney volume and total kidney volume (excluding cysts > 0.2mm).

### CRISPR/Cas9 targeting *ttc30a* replicates the phenotype of *ift80* and *ift172* targeted embryos

We designed sgRNAs targeting all *Xenopus* orthologs of the four candidates for CRISPR/Cas9 mediated loss-of-function experiments (Fig. 4 b, c; Supplement Fig. 4 a-d). Interestingly, *cfap20* and *ttc30a* crispants formed edema suggestive of embryonic renal excretion defects. This effect was specific as co-injection with corresponding mRNAs significantly rescued the edema phenotype (Figure 4 c, Suppl. Fig. 4 c, d).

In order to evaluate a potential effect on limb development, we raised *cfap20*, *ttc30a*, *c11orf74*, and *cep192* CRISPR targeted tadpoles to metamorphosis. *cep192* LOF resulted in a slowed development compared to controls and therefore retained a small stature, while *c11orf74* and *cfap20* crispants did not show any obvious phenotypes and limb formation was normal in all three conditions (Suppl. Fig. 4 j, k). Froglets with *ttc30a* mutations, however, developed severe limb defects (Fig. 4 d, green arrowheads). Extremities were generally shorter and thickened, digits were shortened and demonstrated a polydactyly phenotype (Fig. 4 d; Suppl. Fig. 4 h). MicroCT scans found large depositions of cartilage. Histology revealed a chondrocyte differentiation defect, as previously observed for *ift80* and *ift172* targeted animals. *Ttc30a* targeted embryos had reduced fore- and hindlimb length (Fig. 4 d) and analysis of the kidneys revealed a significantly higher number of cysts and in the total cyst volume (normalized to kidney size) on the injected side of the froglets (Fig. 4 e). Excretion assays confirmed that renal tubular uptake of dextran injection into the coelom was drastically reduced (Fig. 5 a, b), and epidermal cilia were shorter or absent in embryos injected with sgRNA and RNPs targeting *ttc30a* (Fig. 5 c, d). In conclusion, CRISPR targeting *ttc30a* in *Xenopus* replicated all phenotypes observed when targeting the chondrodysplasia genes *ift80* and *ift172*.

**Figure 5:**
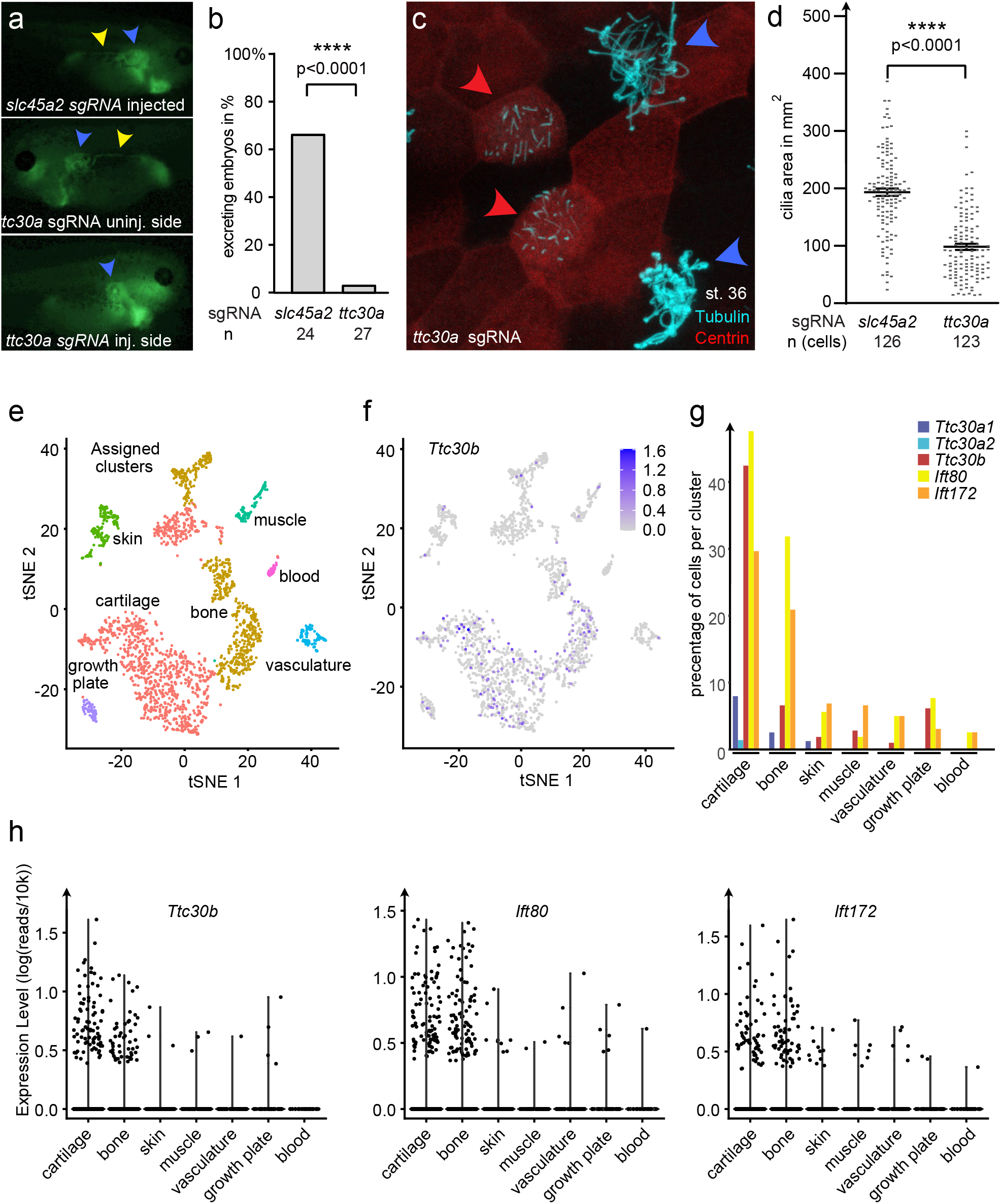
scRNA-Seq data analysis of *ttc30a/b*-positive cells in limbs. (a, b) Excretion assay of *slc45a2* sgRNA injected controls and *ttc30a* targeted embryos. Blue arrowheads point to the proximal tubule, yellow arrowheads highlight fluorescent dextran in the distal tubules as a measure of excretion. (c) Confocal images of epidermal multiciliated cells. centrin-RFP (red) marks cells targeted for *ttc30a* (red arrowheads). Blue arrowheads label wild-type cells. Cilia are stained with acetylated gamma-tubulin (cyan). (d) Quantification of ciliated area per cell in *slc45a2* and *ttc30a* CRISPR targeted cells. Error bars indicate SEM. p>0.05 ns (not significant); p<0.05 *; p<0.01 **; p<0.001 *** (e) Clustering of scRNA-Seq data from mice limbs of E15.5 into 10 different cell clusters using typical marker genes. (f) Detection of *Ttc30b* expression in respective cell clusters. (g) Comparison of percentage of cells per cluster expressing *Ttc30a1*, *Ttc30a2*, *Ttc30b*, *Ift80* and *Ift172*. (H) Expression level of *Ttc30b*, *Ift80* and *Ift172* in respective tissue clusters.

To elucidate the expression pattern of mammalian orthologs of *ttc30a*, we analysed publicly available single cell sequencing data of E13.5 mouse limb buds (Suppl. Fig. 5)^93^. We found that among the three orthologs, *Ttc30b* and to a lesser degree *Ttc30a1* were most prominently expressed in cartilage, bone and growth plate clusters (Fig. 5 e-g). The distribution of cell proportions expressing *Ttc30b* was similar to that of *Ift80* and *Ift172* (Fig. 5 g, h). Thus, the expression pattern of *Ttc30b* and *Ttc30a1* suggest a potential role in cartilage formation of mammals. We screened for *TTC30A/B* mutations in a cohort of 24 unresolved severe ciliary chondrodysplasia cases by exome sequencing, but no likely pathogenic variants were identified. Screening of whole exome sequencing data in a worldwide cohort of more than 500 patients with nephronophthisis related ciliopathies and cystic kidney disease also did not reveal potentially causative recessive mutations in either *TTC30A* or *TTC30B*.

## Discussion

Skeletal ciliopathies are caused by pathogenic mutations in genes that encode ciliary proteins. Thus, Mendelian genetics put the spotlight on the primary cilium as a central organelle in the pathophysiology of chondrodysplasias. This led to the insight that several clinically distinct disease entities may have a common molecular origin and are now recognized as an united disease group4. Today, a causative mutation can be found in almost all patients with SC, both in frequently affected genes and rare loci, which may only occur in a few patients worldwide.

Despite these enormous achievements, a full understanding of all disease-associated proteins, their functional relationships and their molecular functions is incomplete. Therefore, novel *in vivo* models can help to assign tissue specific functions of ciliary proteins molecularly characterized *in vitro*. The tetrapod *Xenopus tropicalis* is both easy to manipulate but not burdened by genome duplications that complicate genetic analysis in other aquatic models^94,95^. Thus, our work demonstrates that disruption of *Xenopus* orthologs of human SC disease genes elicit phenotypes highly consistent with the clinical presentation in patients.

Early limb bud development in *Xenopus* is very similar to that of other tetrapods on a molecular level^52^. As in humans, mouse, and chicken, cilia and cilia-related signaling pathways (Hedgehog (Hh) and Wnt) appear to play an essential role in shaping cartilage and future bone size also in *Xenopus*. Most SC-genes have been described to influence Hh signaling^4^. Among the Hh-ligands, Sonic hedgehog (Shh) defines the anterior-posterior axis of the limb and therefore the number of digits^96^, while Indian hedgehog (Ihh) forms a gradient in the growth plate which controls chondrocyte proliferation versus maturation and therefore bone lengthening^35,97^. As we observed phenotypes consistent with defects in Shh (polydactyly) and Ihh (brachydactyly), both pathways may be affected in our model^35,36,98^. Interestingly, knockout mice of a cilia-dependent regulator of both ligands, *gpr161*, shows the same phenotype, which is however limited to the forelimbs^99^.

Wnt signaling has multiple functions during limb development, but it is not known to what degree Wnt signaling is affected in SC patients^100^. It has been described that Wnt/β-Catenin signaling, in concert with Shh, acts in chondrocyte differentiation and is balanced by ift80^48^. Some SC-genes are linked to Wnt/PCP (planar cell polarity) signaling, which affects the organization of chondrocytes^26,27,101,102^. Indeed, polysyndactyly in combination with brachydactyly, that we also observe in our SC-models, is typical for a loss of *HOXD13*, a gene that regulates *Shh* as well as *WNT5a* (Wnt/PCP) during limb development^101,103,104^. A potential involvement of both, Wnt and Hh pathways may explain why we detected changes in transcripts that impact with both signaling pathways during embryogenesis (Fig. 3 a).

A trove of existing datasets, proteomic and functional screens have contributed to a better understanding of the protein composition of cilia^105,106^. Our data-mining approach integrated these high quality datasets to provide underappreciated links between ciliary proteins and human disease, and ciliopathy subtypes in particular. We created a web application to make the resource lists used in our *in silico* screen accessible and searchable (http://dormantdata.org). Adding further datasets and using novel machine learning algorithms may further enhance the power of a data-driven discovery process. Such refinements could make this approach be applicable to other non-ciliary diseases in the future. By combining the *in silico* screen with *in vivo* modelling, we identified *Xenopus* Ttc30a as a protein involved in chondrocyte differentiation and renal function. It is likely that not all SC relevant components are known. Potentially deleterious variants may be too rare to be identified by Mendelian genetics because the natural mutational load does not reach saturation for all possible pathogenic mutations to occur, or lead to early embryonic lethality. In addition, essential proteins may be protected by genetic compensation of functionally redundant genes^107,108^. The human *TTC30A/B* locus may represent a critical site that is protected by early and reoccurring duplication events in the tetrapod phylogeny that increasingly relied on limbs for locomotion and dexterity (Fig. 4 a). TTC30A/B are highly conserved in eukaryotes. The amino acid sequence between the green algae *Chlamydomonas rheinhardtii* and human TTC30A is 73% identical^90^. In zebrafish, the *fleer* (*ttc30a*) mutant shows a general ciliopathy phenotype, including pronephric cysts, dorso-ventral body axis curvature, laterality defects, hydrocephalus, and retinal degeneration^91,109^. Cilia are either shorter or absent (pronephros, Kupffer’s vesicle and olfactory placode) and lack parts of the microtubule-duplets of the axoneme. Both, *in vitro* and *in vivo* experiments identified ttc30a as a peripheral protein of the IFT-B1 subcomplex^110,111^, where it links the ift88/ift52-complex with Kif-proteins (Kif17)^112^. Further, ttc30a is necessary for polyglutamylation and polyglycylation of tubulin by regulating the ciliary activity of ttll3/6 (Tubulin Tyrosine Ligase Like 3/ 6), and ccp5 (cytoplasmic carboxypeptide 5)^15,92^. Both posttranslational modifications stabilize tubulin complexes and are necessary for cilia motility^92^. Interestingly polyglutamylation regulates velocity of the Kinesin-2 complex and therefore the uptake of Hedgehog components (Smo, Gli3) after signaling activation, and has recently been found to be dependent of the Joubert syndrome associated proteins ARL13B and CEP41^18,19,113^. It is not known whether tubulin polyglutamylation is affected in other ciliopathies. Intriguingly, the ciliary localization of PKD2 was found to depend on tubulin glutamylation, suggesting a role in renal cystogenesis^18,114^.

Like glutamylation, tubulin glycylation appears to be present only in primary cilia of certain cell types. Glycylation was detected in MDCK cells, but absent in IMCDs and RPEs^16^. Only a subset of cilia in the spinal cord and pronephros are glycylated, but not Kupffer’s vesicle cilia of zebrafish^92^. Thus, tubulin amino acid PTMs not only constitute a part of the “tubulin code” that diversifies ciliary functions, but may contribute to the pleiotropic phenotypes of ciliopathies^114^.

The molecular function of TTC30A/B in ciliogenesis is well established, but has not been associated with skeletal defects. In human tissue, TTC30A and TTC30B are strongly expressed in ciliated tissues, such as airway epithelial cells, ciliated cells of the ductuli efferentes in the epididymis, and the spermatozoa progenitor tissue of the seminiferous ducts^115^. Consistent with our own analysis, single cell RNA-Seq data of embryonic mice revealed an enhanced expression of *ttc30a1*, *ttc30a2* and *ttc30b* in osteoblasts and chondrocytes (Fig. 5)^116^. A rare missense variant in *TTC30B* (rc.1157C > T, p.A375V) has recently been described in a large Chinese family with polysyndactyly, suggesting an autosomal dominant mode of inheritance^117^. However, no other unifying clinical phenotypes were observed in the 27 affected family members. *In vitro* analysis suggested that downregulation of *TTC30B* had an inhibitory effect on Shh-pathway stimulation, consistent with our findings that Ttc30a has a role in limb patterning. Hedgehog signalling does not influence cystogenesis in autosomal polycystic kidney disease (ADPKD), suggesting that other ciliary mechanisms are more likely to be at play^118^. Further investigation will need to elucidate, which signals are disrupted by loss of Ttc30a in renal cyst formation and to what degree polyglutamylation and - glycylation are more widely implicated in nephronophthisis and other ciliopathies.

## Material and Methods

### Animal maintenance and handling

Adult animals were kept according to the German and Swiss law for care and handling of research animals. Husbandry and treatment were approved by the local authorities (Regierungspräsidium Freiburg and Veterinäramt Zürich).

*Xenopus* embryos were kept in 0.3x Marc’s Modified Ringer (MMR, HEPES (free acid) 5 mM, EDT 0.1 mM, NaCl 100 mM, KCl 2 mM, MgCl_2_ 1 mM, CaCl_2_ 2 mM) with 0.05 mg/ml Gentamycin. Pre-metamorphic tadpoles were raised on Sea Micron (Heinsberg, Germany) at room temperature with daily buffer/water exchange. After approximately 4 weeks the food was changed to Sera Vipagran Baby (Heinsberg, Germany) and after approx. 8 weeks crushed adult frog food (Shrimp Sticks) was fed. Staging was performed according to Nieuwkoop and Faber^119^.

### Plasmids, mRNAs, sgRNAs and MOs

For gene expression analysis of SC reference and candidate genes, *X. laevis* gene variants were either cloned into pGEM-T (*iIft80*) or pBluescript (*iIft172*) or a PCR-product was used as template for antisense probe synthesis (*ttc30a) (X. laevis*) after addition of the T7 promoter site to the reverse primer.

The constructs *pax2-a* and *pax6* (*X. laevis*) were previously cloned in our lab^120^. *prox1*, *aplnr*, *nkx2.5*, *gli2*, *patched1*, *cacgn1*, *ccng1*, *cfh*, *olfm4*, *ulk1* and *riok3* (*X. tropicalis*) were cloned into pGEM-T. Further constructs were kindly provided by M. Blum (*snail*^121^, *twist1.S*^122^ and *sox9*), J.B. Wallingford (*nkx2.2*)^123^, N. Ueno (*nkcc2*) and O. Wessely (*sglt1k*)^124^, all *X. laevis*. For antisense probe preparation, plasmids were linearized and transcribed with T3, T7, or SP6 polymerase (Roche, Basel, Switzerland). The coding sequence of *X. tropicalis ift80*, *ift172*, and *ttc30a* were cloned into VF10^6^ for rescue experiments. Constructs were linearized with SalI (*ift172*, *X. laevis* and *ttc30a, X. tropicalis*) or AscI (*ift80*, *X. laevis*) for mRNA synthesis. mRNA was made with mMESSAGE mMACHINE T7 Transcription Kit (Invitrogen, AM1344) and cleaned up with RNeasy mini kit (Qiagen, Cat No./ID: 74104).

Short guide RNAs were designed with chopchop and CRISPRscan (https://chopchop.cbu.uib.no; https://www.crisprscan.org) for CRISPR/Cas9 targeting^125,126^. The PCR-based method was used for sgRNA synthesis described by Nakayama et al. 2013^127^. The PCR-product was amplified with the KOD polymerase (Novagen; 71086) and purified with the QIAquick PCR Purification Kit (Qiagen; Cat No./ID: 28104). sgRNAs were synthesized overnight with the MEGAscript T7 Transcription Kit (Invitrogen; AM1334). Afterwards sgRNAs were cleaned-up with the mirVana miRNA Isolation Kit (Ambion; AM1560). Several sgRNAs were tested for each gene and the most efficient sgRNA was used for experiments, as determined by ICE or HRMA analysis. For *X. laevis* experiments, previously published tyrosinase sgRNAs was used^53^.

To verify functionality of *ift80* and *ift172* sgRNA in experiments where unilateral injections for limb analysis were performed, high dose injections at 1-cell stages were done in parallel, and edema formation was confirmed to occur at stages 42-45 (Suppl. Fig. 1n, 4g).

Morpholino oligonucleotides for *X. laevis Ift80*, *Ift172* and *Ift52* were designed by and ordered from genetools. Additionally the Standard Control MO from genetools was used for injected controls. Sequences of primer and morpholino oligonucleotides are given in Supp. Table 5.

### Microinjections

Embryos were obtained by *in vitro* fertilization (*X. laevis* and *tropicalis*) or natural mating (*X. tropicalis*). For injections, embryos were transferred to 3% Ficoll with 0.1% BSA (*X. tropicalis*) or 2% Ficoll (*X. laevis*) dissolved in 0.3x MMR. Depending on the experiment embryos were injected at the 1-cell stage (phenotype analysis, genotyping and RNA-Seq) or unilaterally at 2-4 cell stage (all other experiments). The injected volume was 10nl in *X. laevis* and 5nl in *X. tropicalis*. In all experiments a lineage tracer (2ng Fluorescein-Dextran 70kMW - Invitrogen; D1823) was co-injected, and embryos sorted according to the injected side at neurula stage. For experiments on epidermal cilia, 0.2ng Centrin-mRFP (kindly provided by J.B. Wallingford)^128^ was co-injected.

For CRISPR/Cas9 experiments, 0.175ng sgRNA was injected in combination with 0.6ng Cas9 protein with NLS (PNABio; CP01). As a control, sgRNA against *slc45a2* (*X. tropicalis*) or tyrosinase (*X. laevis*) was used. Rescue experiments were performed with 0.1ng *ift80*, 0.15ng *iIft172* or 0.2ng *ttc30a X. tropicalis* mRNA. For raising crispants 0.015-0.02ng *ift80* or *ift172*-sgRNAs, 0.02-0.05ng, and *ttc30a* sgRNA was injected at the 2-cell stage. The same amount was used for controls.

In *X. laevis* 3.6 - 14.5ng *ift80* TB MO; 3.6 - 7.25ng *ift172* TB MO as well as *ift52* TB MO were injected in knockdown experiments. In *X. tropicalis* half of the amount was used. The same concentration of a Standard Control MO was injected as control.

### Genotyping

sgRNA efficiency was tested by genotyping embryos after 1 to 3 days. Five injected individuals per sgRNA were lysed and DNA extracted with the DNeasy Blood and Tissue Kit (Qiagen; Cat No./ID 69504). Genotyping was performed with high resolution melting analysis^129^ using the MeltDoctor™ HRM Master Mix (Applied Biosytems) on a Roche LightCycler 480. The DNA-product of the analysis was purified (QIAquick PCR Purification Kit) and Sanger sequenced. Sequencing results were analyzed with ApE (A plasmid Editor v2.0.50b3) and the ICE-tool from Synthego (https://ice.synthego.com/#/).

### Transcriptomic analysis

Three independent replicates of *ift80* and *ift172* CRISPR/Cas9 targeting experiments were analyzed by RNA-seq. One-cell stage embryos were injected, and Cas9 protein injections served as negative controls. Embryos were lysed at stage 22 and RNA isolated with the RNeasy mini kit (Qiagen, Cat No./ID: 74104). Afterwards, genomic DNA was extracted from the columns using 8mM NaOH on the filter to solve the DNA and EDTA (0.57 mM) and HEPES (426 mM) to stabilize it. HRMA and ICE analysis were performed to confirm successful indel formation. RNA was further purified with RNeasy Universal Plus kit from Qiagen (Cat No./ID: 73404) and Illumina sequenced by GATC (Germany).

RNA-seq analysis was performed on the Galaxy platform^130^. Raw reads were trimmed (Trim Galore!) and aligned to the *X. tropicalis* genome version 9.1 using STAR^131^. Gene counts were calculated using featureCounts and differential expression was determined by DESeq2, taking batch effects of the replicates into account^131,132^. For single cell RNA analysis, disassociated limbs data for E15.5 time point from GEO accession number GSE142425 was analysed using R v4.0.2 and Seurat v3.1.5^133,134^. Cells expressing less than 1200 or more than 4200 genes and a mitochondrial gene percentage of >10% were excluded in QC leaving 2142 cells in total. Highly variable genes (N=2500) were extracted from the normalized data, scaled and further used in downstream analysis. Data was clustered with t-distributed stochastic neighbor embedding (tSNE) using first 10 PCs for dimensionality. Cell types were assigned for 10 clusters (eliminating three clusters) using the markers reported in Kelly et al.^93^.

### Analysis of craniofacial defects

For early phenotype analysis, embryos were fixed with MEMFA (0.1 mol/L MOPS (3-morpholinopropane-1-sulfonic acid), 2 mmol/L EGTA, 1 mmol/L MgSO_4_, 3.7% formaldehyde, pH 7.4) at stage 42 – 45 for approx. 2h at room temperature and transferred to 1x phosphate-buffered saline (PBS) for imaging. Pictures were taken from the dorsal side of the embryo with a SteREO Discovery.V8 microscope and ZEN 2011 (blue edition; Zeiss, Oberkochen, Germany) software. The shortest distance between the eyes was measured using the line tool of imageJ 2.0.0. As analyzed embryos of different batches differed in age, the measured distances were normalized to the control embryos of the same batch.

### Whole mount *in situ* hybridization

Whole mount *in situ* hybridization was performed as described by Sive et al.^135^. Embryos were fixed at desired stages with MEMFA for 1.5-2h at room temperature. Antisense probes were detected with a phosphatase-conjugated secondary DIG-antibody (Roche, Basel, Switzerland). Stained embryos were bleached for 1-2h with H_2_O_2_ and methanol (1:2). Afterwards they were rehydrated with 1x PBS and imaged with a SteREO Discovery.V8 microscope and ZEN 2011 (blue edition) software.

### Immunostaining

For immunostaining, embryos were fixed for 1-2h in MEMFA. The staining was performed as described before^136^. Cilia were detected with a monoclonal anti-acetylated tubulin antibody (Sigma, T7451) in combination with an anti-mouse secondary Alexa Fluor 488 antibody (Invitrogen, A-11001). Pictures were taken with a ZEISS LSM 510 DUO with inverted microscope Axiovert 200 and a 63x LCl-Plan Neofluar objective and a Leica SP8 inverse FALCON and a 63x HC PL APO CS2 objective. Ciliation was determined in Centrin-RFP-positive cells and the area of their acetylated tubulin staining was measured on maximum intensity projections with ImageJ 2.0.0.

For pronephros analysis a fluorescein–labeled *Lycopersicon esculentum* lectin (1:100 dilution; Vector Laboratories) was used for visualization. A SteREO Discovery.V8 from Zeiss and Zen2011 Blue Edition was used for imaging. Pronephros size was measured with the line tool of ImageJ 2.0.0 and the ratio of injected versus uninjected side calculated^6^.

### Excretion assay

Excretion assays were performed as described before^137^. Embryos were anesthetized with 0.2mg/ml MS-222 around stage 38. Approximately 20nl of Fluorescein-labeled Dextran (70kMW - Invitrogen; D1823) was injected into the coelomic cavity around the heart. Uptake and excretion of Dextran was documented by performing time-lapse microscopy (2 fps for a time-span of 4 minutes). Pictures were taken with the SPOT Insight FireWire system (Diagnostic Instruments) on a Leica MZ16 stereomicroscope and the SPOT advanced software. For visualization time-lapse movies were converted to AVI-movies of 40fps with imageJ 2.0.0.

### Analysis of CRISPR/Cas9 froglets

Froglets for limb analysis were euthanized and fixed in MEMFA. Iodine contrast enhancement was done by washing froglets in 1x PBS Buffer for 2 days, followed by dehydration to ethanol, and staining in 1% iodine solution (Sigma-Aldrich, 207772) in ethanol. The day before microCT scans, froglets were washed with 100% ethanol overnight. The animals were scanned with a Caliper Quantum Fx from PerkinElmer. To stabilize froglets in the microCT scanner, a piece of 1% agarose solved in water and dehydrated in 100% ethanol was used. MicroCT DICOM pictures were converted with Imaris File Converter 9.5.1 to imaris files. Imaris 9.5.1 was used to determine the volume of kidneys and the volume of kidney cysts. Length of limbs was measured with RadiAnt DICOM Viewer 5.5.1. To determine the total length of the hindlimb, length of femur and tibia/fibula was measured and summed. Length of forelimbs was determined by adding the length of humerus and ulna/radius.

### Histology

Iodine staining of froglets was removed with Thiosulfate as described in Hopkins et al. 2015^138^. Skin and muscles were removed from limbs and bone/cartilage washed with PBS. Preparations were decalcified with 20% EDTA - citric acid, pH 7.5 solution (BIOCYC, 400500201) for 3-5 days at room temperature. Limbs were dehydrated and embedded in paraffin. 5μm sections were made with the rotary microtome Microm HM355S. Sections were stained with Hematoxylin and Eosin following standard methods.

### *In silico* screen to identify novel SC-candiates

For the *in silico* screen, we collected 90 publicly available datasets representing the results of various screens and unbiased large-scale experiments in various species (resource lists, Suppl Table 3). Gene names were mapped to the human orthologue (HGNC) using biomart and manually curated. Online Mendelian Inheritance in Man (OMIM) “phenotypic series” data was aggregated from the genemap.txt file downloaded on 26.01.2017 to obtain the 388 input lists of known and confirmed genetic diseases by matching phenotype descriptions to gene symbols.

First, each resource list was tested by Fisher’s exact test, in which multiple testing was accounted for by use of a Bonferroni correction for the number or resource lists, to determine if the genes in the input list were enriched (p<0.05/90) in a given resource list^139^. Genes on resource lists that showed significant enrichment of known monogenic genes on input lists were scored according to either their position in ranked resource lists 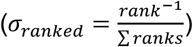 or by the total number of genes in the resource list 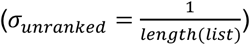. Then, combined scores across all enriched resource lists were computed.

Finally, we drew 10 million random input lists and scored genes analogously to estimate the background distribution of scores, compute resampling based enrichment p-values and to adjust for multiple testing by the Benjamini Hochberg procedure (FDR <0.05).

### Direct Mutation analysis of chondrodysplasia patients

DNA was extracted by standard protocols from fetal fibroblasts and FFPE (Formalin-fixed paraffin-embedded) tissue (Qiagen). PCR-amplifications of exons and intron-exon boundaries of TTC30A (ENST00000355689.6) and *TTC30B* (ENST00000408939.4) were analyzed with an ABI3130xl capillary sequencer (Life Technologies) to detect putative mutations^140,141^. Allele frequency was bioinformatically analyzed using an in-house exome database, the 1000 Genomes Project (http://www.internationalgenome.org/), the NHLBI Exome Sequencing Project (http://evs.gs.washington.edu/EVS/), and the dbSNP database (https://www.ncbi.nlm.nih.gov/projects/SNP/).

### Whole exome sequencing of chondrodysplasia patients

The SureSelect Exome Enrichment kit V7 (Agilent Technologies) was used to enrich exome sequences. DNA was sequenced with a NextSeq 550 System (Illumina) resulting in 2×150-bp paired-end reads. A workflow based on the Genome Analysis Toolkit best practice was used to analyze the sequencing results bioinformatically^142^. For annotation and further processing the Ensembl Variant Effect Predictor software and GEMININ were used^143–145^.

### Whole exome sequencing of nephronophthisis patients

Following informed consent, we obtained clinical data, pedigree data, and blood samples from individuals with nephronophthisis-related ciliopathies (NPHP-RC) from worldwide sources using a standardized questionnaire. Approval for human subject research was obtained from the Institutional Review Boards of the University of Michigan and Boston Children’s Hospital. Informed consent was obtained from the individuals and/or legal guardians, as appropriate. The diagnosis of NPHP-RC was made by nephrologists based on relevant imaging.

Whole exome sequencing (WES) was performed as previously described^146^. Briefly, DNA samples from affected individuals and unaffected family members were subjected to WES using Agilent SureSelect™ human exome capture arrays (Life Technologies) with next generation sequencing (NGS) on an Illumina™ sequencing platform. Sequence reads were mapped against the human reference genome (NCBI build 37/hg19) using CLC Genomics Workbench (version 5.0.1) software (QIAGEN). Mutation analysis was performed under recessive, dominant or de novo models, as previously published^146–148^.

### Statistical analysis

Volume, number of kidney cysts and total kidney volume (excluding cysts) between the injected side of the control (slc45a2) and the injected side of *ift80*, *ift172* or *ttc30a* targeted animals, respectively, were compared with the Mann-Whitney-U test. The same test was used for analysis of eye distance, cilia area measurements, and pronephros length (Lectin) measurements. The paired t-test was used for statistical analysis of limb length. Unpaired t-test statistics were calculated for centrosome number after testing for normal distribution. Differences of edema formation and excretion were analyzed using chi^2^ test. Statistical analysis was performed with GraphPad Prism 8.3.1. (****p<0.0001, ***p<0.001, **p<0.01, and *p<0.05). All measurements were taken from distinct samples. No sample-size calculation was performed before the experiments.

### Data availability statement

The datasets generated during and/or analysed during the current study are available from the corresponding author on reasonable request. The raw data associated with Figure 3a and Supp. Figures 3 c,d was deposited at at SRA with the BioProject accession number: PRJNA670560.

### Code availability statement

The code used in the *in silico* screen is publicly available at https://github.com/genepi-freiburg/GeneSet.

## ACKNOWLEDGEMENTS

We are grateful for technical assistance to Alena Sammarco, Jessika Kleindienst, Marko Vujanovic and Claudia Meyer, and the staff of the Life Imaging Center Freiburg at the Center for Biological Systems Analysis (ZBSA) of the University of Freiburg, the Center for Microscopy and Image Analysis (ZMB) and the Zurich Integrative Rodent Physiology (ZIRP) of the University of Zurich. This study was supported by the German Research Foundation (DFG) to SSL (Emmy Noether Program (Li-Li 1817/2-1), PS (CRC 992) and AK (KO 3598-5/1), and the Swiss National Science Foundation (SNF) to SSL (NCCR Kidney.CH) and (310030_189102/1). FH was supported by the National Institutes of Health (DK_068306).

## AUTHOR CONTRIBUTIONS

S.S.L. and M.G. wrote the manuscript with contributions from A.H., P.S., K.G., K.D., F.Hi., E.L. and A.K.. S.S.L. supervised the project. M.G. performed most experiments and analyzed most of the data. S.S., R.D., F.He., and W.S. contributed to *Xenopus* experiments and data analysis. A.K., A.H., and P.S. performed and analyzed data for the *in silico* screen. K.G. analyzed the RNA-Seq and scRNA-Seq data. F.Hi., K.D. and E.L. screened patient cohorts for TTC30A/TTC30B mutations.

## COMPETING INTERESTS STATEMENT

The authors declare no competing interests.

## Supplementary Figures

**Supplementary Figure 1**: Specificity controls of *ift80* and *ift172* CRISPR/Cas9 targeting experiments

(a) Gene structure of *X. tropicalis ift80* and *ift172* with introns (lines) and exons (boxes). sgRNA-binding-sites are marked by a red arrow. (b, c) CRISPR editing analysis for *ift80* and *ift172*. The site of the expected cut is depicted by black vertical lines in Sanger Sequencing chromatograms, the sgRNA binding site is marked in blue and the PAM site in red. (d) Craniofacial defects of *ift80* and *ift172* CRISPR targeted embryos were analyzed by measuring the distance between the eyes of stage 43-46 *X. tropicalis* tadpoles and normalized to the average of uninjected controls. (e) Normalized eye distances decreased significantly in CRISPR targeted embryos. (f) Knockdown of *ift80* and *ift172* using anitsense morpholino oligonucleotides. Knockdown resulted in edema formation. (g, h) Decrease of eye-distance for *ift80* and *ift172* CRISPR targeted *X. laevis* embryos. (j) CRISPR targeting of *ift80* and *ift172* in *X. laevis* leads to edema formation, (k) *In situ* hybridization detected expression of *ift80* and *ift172* in limb buds of *X. laevis* (black arrowhead). po - posterior; a - anterior; pr - proximal; d - distal (l) Categorization of mutant froglets according to phenotypic strength of limb malformations.

Error bars indicate SEM. p>0.05 ns (not significant); p<0.05 *; p<0.01 **; p<0.001 ***

**Supplementary Figure 2:** Phenotypic analysis of *ift80* and *ift172* targeted embryos.

(a, b) Expression analysis of *ift80* and *ift172* by whole mount *in situ* hybridization in st. 26/27 embryos. The magnified images show a spotty pattern on the epidermis. (c, d) Expression analysis of *aplnr*, *nkx2.5*, *prox1*, *nkcc2* and *sglt1* of *X. tropicalis* (c) and *X. laevis* (d) unilaterally CRISPR targeted tadpoles. The bar graphs indicate a stronger (gray) or weaker (black) signal on the injected side.

(e, f) Tomato-Lectin stain visualizes the pronephric tubules in unilaterally CRISPR targeted stage 39 tadpoles. Measurements of the bounding box area of the proximal tubule was not different between the injected side of the embryo and the wildtype half in each case. (g) Quantification of centrioles of epidermal MCCs after injection of *centrin-RFP* mRNA together with sgRNAs and Cas9. There was no significant difference in centriole number in CRISPR targeted cells. Error bars indicate SEM. p>0.05 ns (not significant); p<0.05 *; p<0.01 **; p<0.001 ***

**Supplementary Figure 3:** RNA-Seq analysis of *ift80* and *ift172* targeted embryos.

(a) Schematic representation of the RNA and DNA extraction protocol used for RN-Seq experiments. Embryos were genotyped before sending purified RNA for sequencing (see “materials and methods” for detailed description).

(b) Indel- and knockout scores from ICE for experimental replicates used in the RNA-Seq experiment. (c, d) Vulcano-plots of RNA-Seq analysis of *ift80* and *ift172* targeted embryos show that the targeted genes were downregulated efficiently in the respective experiments (red arrowheads). (e) *In situ* hybridization for differentially expressed genes in the RNA-Seq screen. *Col1a1* expression is shown for *X. laevis*, all others for *X. tropicalis* tadpoles. (f, g) *In situ* hybridization of tissue marker genes was examined in *X. tropicalis* (f) and *X. laevis* (g) after unilateral CRISPR targeting of *ift80* and *ift172*. The quantifications indicate a stronger (gray) or weaker (black) signal on the injected side. No obvious changes were found, only the *snail* expression, marking cranial neural crest streams, appeared to be fused in *ift80* and *ift172* targeted embryos.

**Supplementary Figure 4:** Analysis of SC-associated candidate genes

(a) Gene structure of *X. tropicalis ttc30a* with introns as lines and exons as boxes. The sgRNA target site is marked by a red arrow. (b) Genotyping and ICE analysis of *ttc30a* targeted embryos (sgRNA binding site - blue; PAM - red; cutting site - vertical black line). (c, d) Percentage of embryos developing edema after targeting *c11orf74*, *cfap20* and *cep192*. Targeting of *cfap20* resulted in significant edema formation, and was rescued by co-injection of *cfap20* mRNA. Targeting *cep192* resulted in significant edema formation, but at low percentages. (e, f) Craniofacial analysis revealed that the distance between the eyes was smaller for *cep192*, *ttc30a*, and *cfap20* targeted embryos, but not *c11orf74*. (g) Percentage of edema formation in embryos injected at the one cell state in parallel to the animals raised to the froglet stage were used as efficacy control. (h) An example of a *ttc30a* mutant froglet with polydactyly (orange arrowheads) and both fore- and hindlimb malformations (red arrowheads). Embryos were injected at the 2-cell stage into the right blastomere with sgRNA targeting *ttc30a* and Cas9. (j) Examples of unilaterally *cep192* and *cfap20* targeted animals without an obvious phenotype. (k) Quantification of limb defects of unilaterally targeted froglets. Shortened limbs were only detected in *ttc30a* targeted animals. Error bars indicate SEM. p>0.05 ns (not significant); p<0.05 *; p<0.01 **; p<0.001 ***

**Supplementary Figure 5:** scRNA-Seq data analysis of embryonic mouse limb buds.

(a) tSNE plot of all identified cell clusters in scRNA-Seq data of E15.5 mice limbs. (b, c) Marker genes used for defining the clusters. Gene expression is indicated in blue on the tSNE-plot. (c) Violin plots of respective marker transcript levels across the assigned tissue clusters. (d) Expression of *Ift80*, *Ift172*, *Ttc30a1* and *Ttc30a2* depicted in blue in the tSNE plot. (e) Expression levels of *Ttc30a1* and *Ttc30a2* in the tissue clusters.

Supplementary Table 1

Differentially regulated transcripts detected by RNA-Seq of *ift80* CRISPR targeted embryos.

Supplementary Table 2

Differentially regulated transcripts detected by RNA-Seq of *ift172* CRISPR targeted embryos.

Supplementary Table 3

Resource List and references used for the *in silico* discovery screen.

Supplementary Table 4

Candidate genes of the *in silico* discovery screen for SRTD at FDR 5%.

Supplementary Table 5

Sequences of primers and morpholino oligonucleotides.

Supplementary Table 6

Results of the *in silico* screen for OMIM diseases/phenotypes at FDR 5%.

(Amyotrophic lateral sclerosis, Bardet-Biedl syndrome, cortical dysplasia, cranioectodermal dysplasia, Diamond-Blackfan anemia, Joubert syndrome, Long-QT syndrome, Meckel syndrome, nephronophthisis, pachyonychia congenita, primary cilia dyskinesia, Senior-Loken syndrome)

Supplementary Movie 1

Fluorescein-dextran was injected into the coelomic cavity and recorded by a timelapse movie (4fps for 3min). AVI-movies are shown with 40fps. Excretion was only observed for *slc45a2* targeted embryos (embryo on top) but not for embryos targeted for SC-genes (*ift80* targeted embryo in the middle, *ift172* targeted embryo at the bottom).

Supplementary Movie 2

Time-lapse movie for excretion in *ttc30a* targeted tadpoles (*slc45a2* targeted embryo on top; *ttc30a* targeted embryos at the bottom).

## Notes

http://dormantdata.org

### Competing Interest Statement

The authors have declared no competing interest.

