## Supplementary material for "CRISPR/Cas9 targeting Ttc30a mimics ciliary chondrodysplasia with polycystic kidney disease": Getwan et al._Supplement.pdf

### Supplementary Figure 1

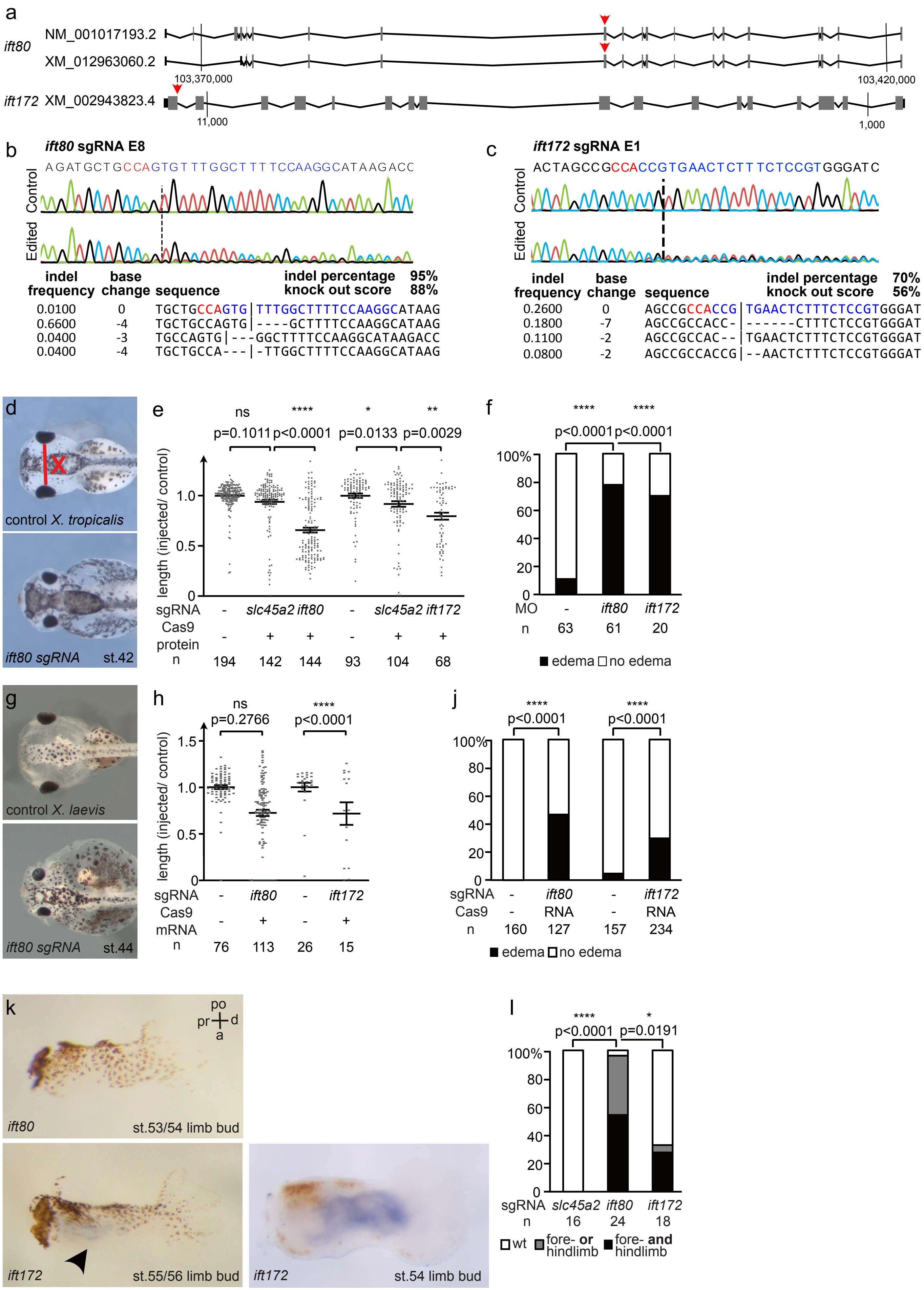

### Supplementary Figure 2

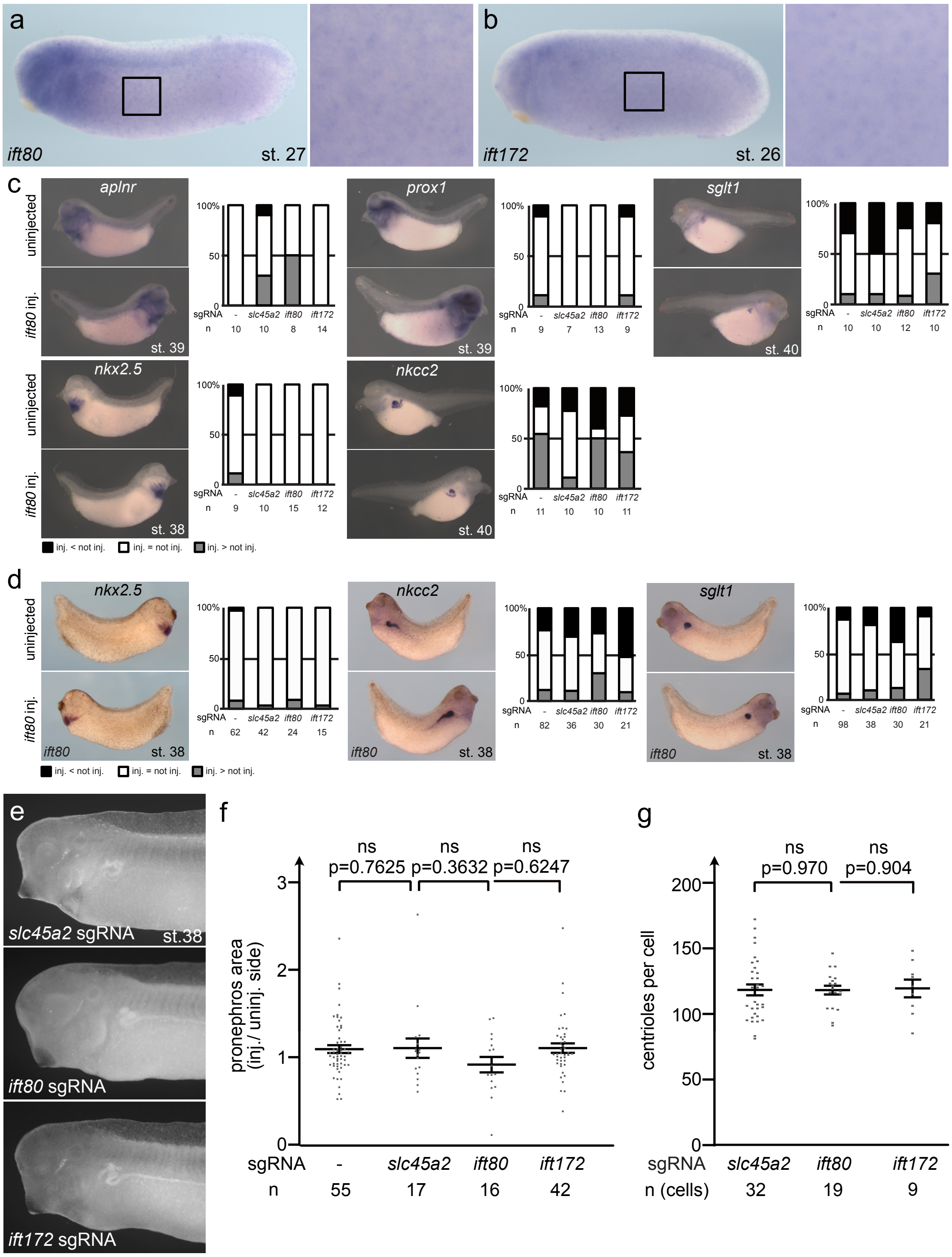

### Supplementary Figure 3

a

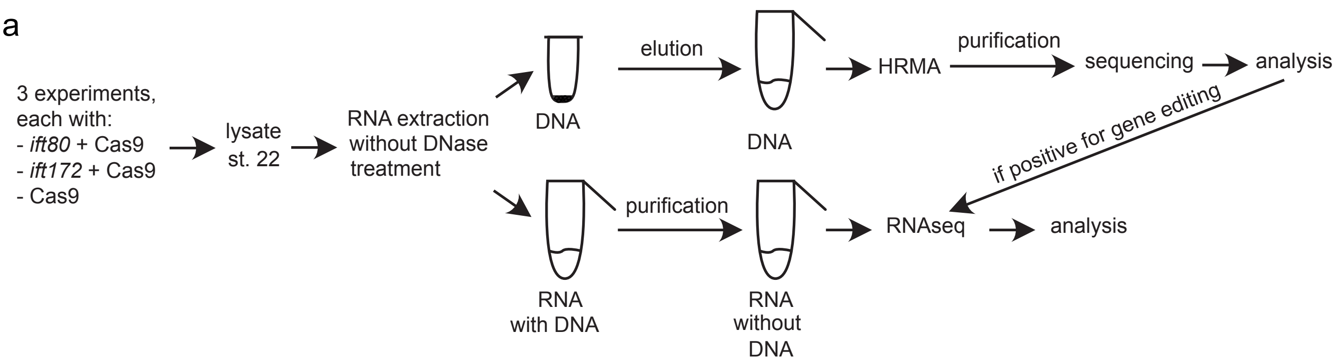


b

| Gene | Experiment | Indels (%) | KO-score (%) |
|---|---|---|---|
| *ift80* | 1 | 56 | 55 |
| *ift80* | 2 | 95 | 88 |
| *ift80* | 3 | 21 | 21 |
| *ift172* | 1 | 17 | 12 |
| *ift172* | 2 | 70 | 56 |
| *ift172* | 3 | 35 | 24 |

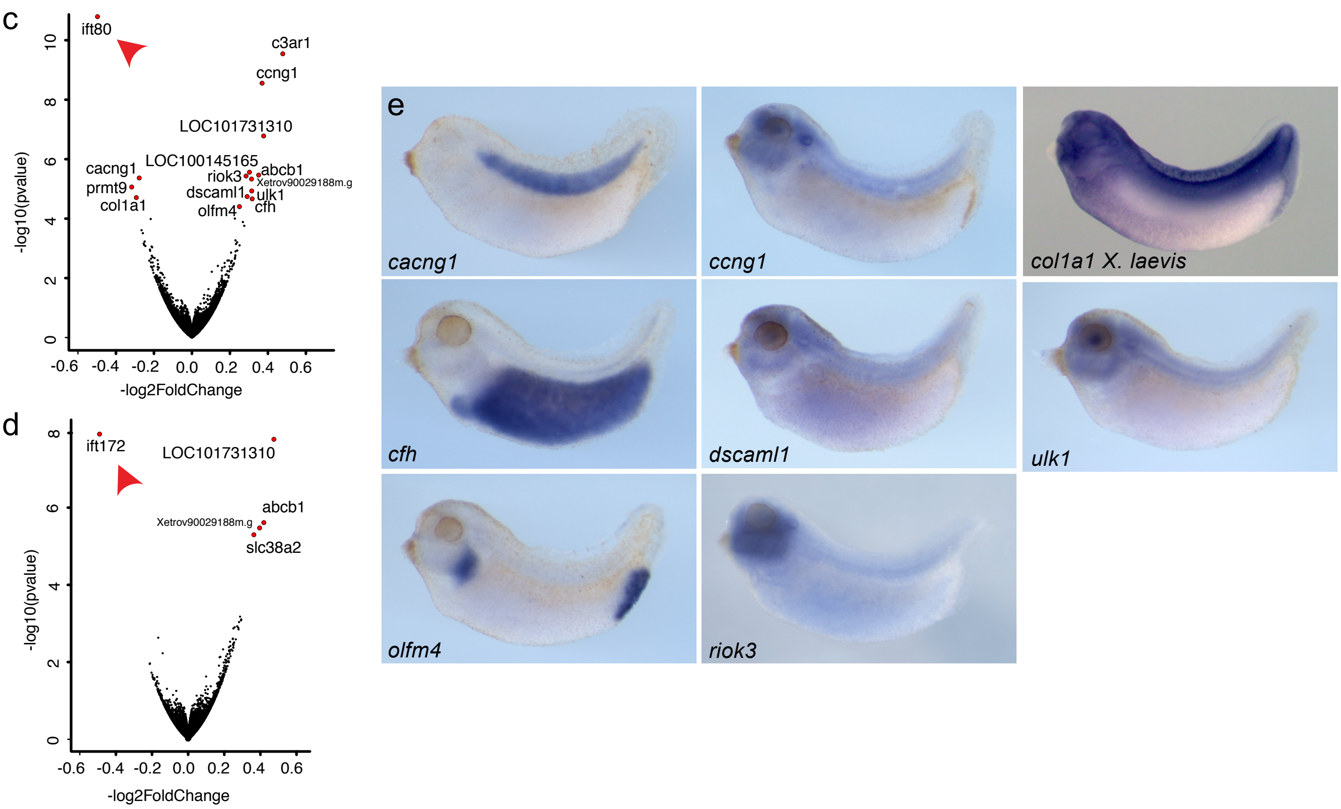


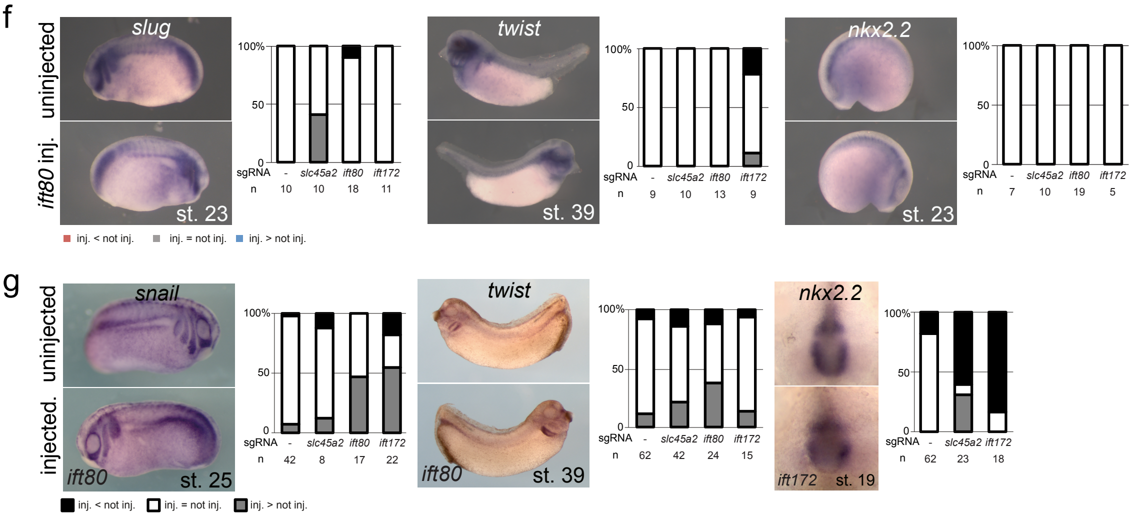

### Supplementary Figure 4

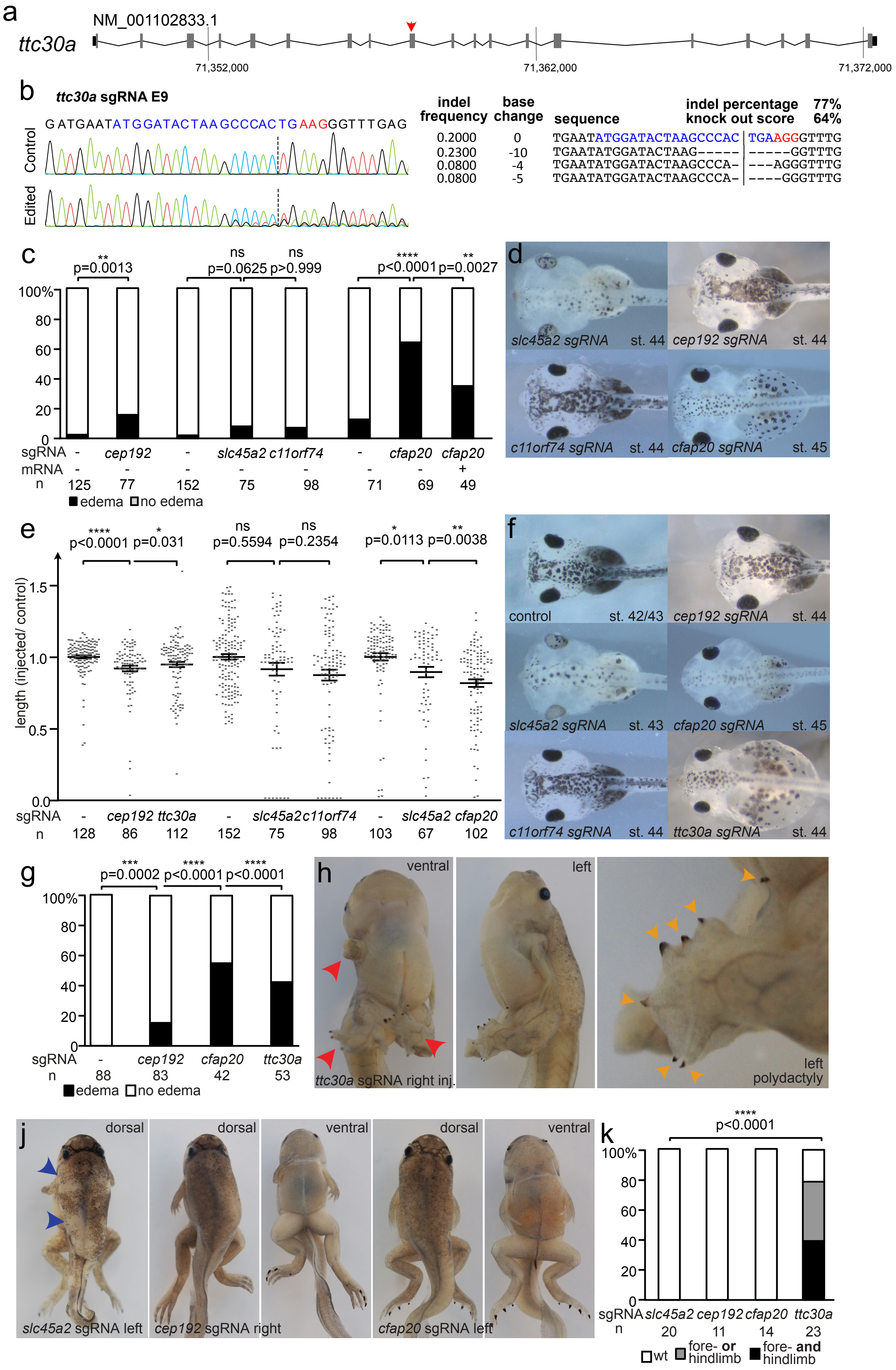

### Supplementary Figure 5

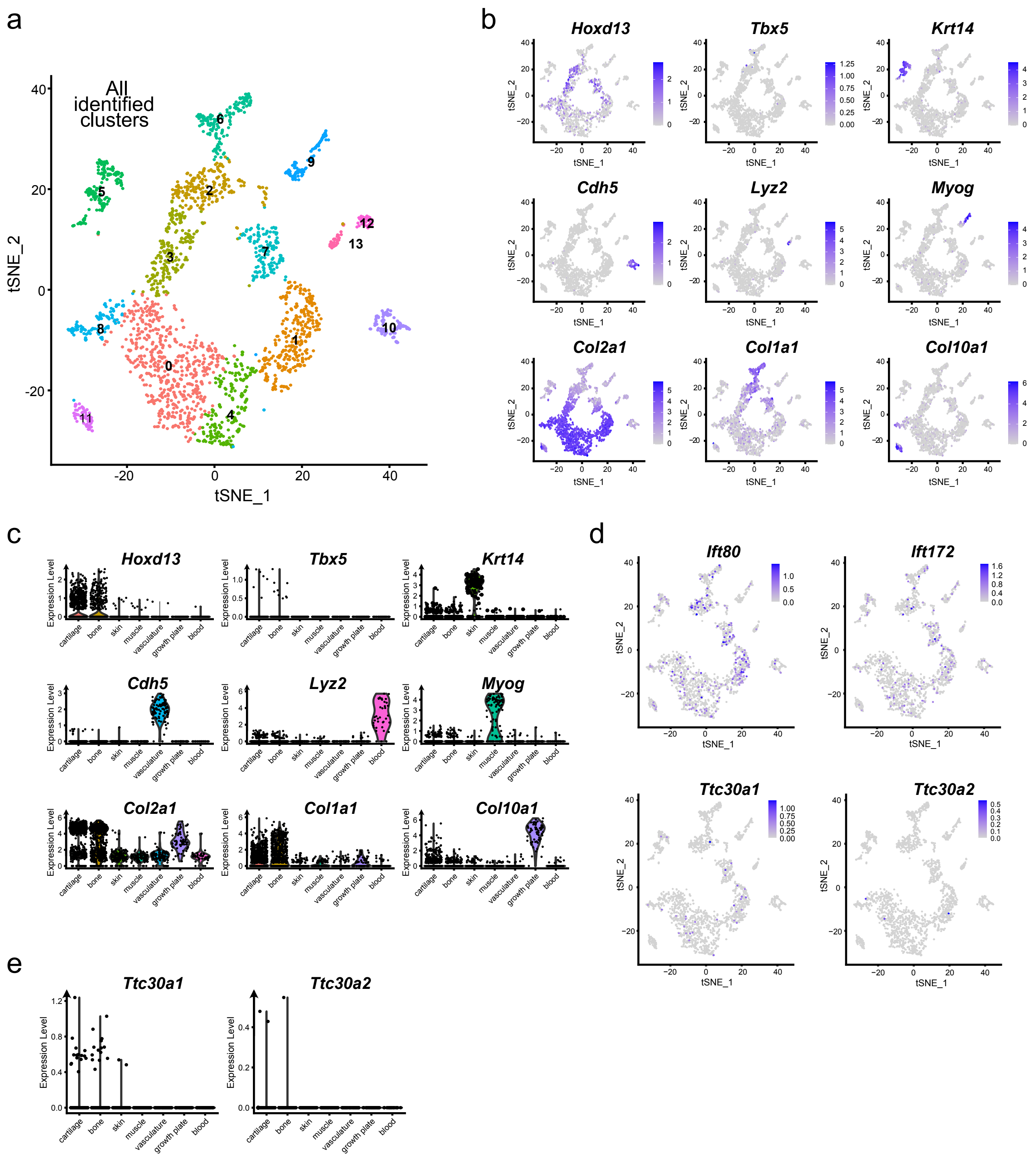

#### Supplementary Figures

**Supplementary Figure 1**: Specificity controls of *ift80* and *ift172* CRISPR/Cas9 targeting experiments
(a) Gene structure of *X. tropicalis ift80* and *ift172* with introns (lines) and exons (boxes). sgRNA-binding-sites are marked by a red arrow. (b, c) CRISPR editing analysis for *ift80* and *ift172*. The site of the expected cut is depicted by black vertical lines in Sanger Sequencing chromatograms, the sgRNA binding site is marked in blue and the PAM site in red. (d) Craniofacial defects of *ift80* and *ift172* CRISPR targeted embryos were analyzed by measuring the distance between the eyes of stage 43-46 *X. tropicalis* tadpoles and normalized to the average of uninjected controls. (e) Normalized eye distances decreased significantly in CRISPR targeted embryos. (f) Knockdown of *ift80* and *ift172* using anitsense morpholino oligonucleotides. Knockdown resulted in edema formation. (g, h) Decrease of eye-distance for *ift80* and *ift172* CRISPR targeted *X. laevis* embryos. (j) CRISPR targeting of *ift80* and *ift172* in *X. laevis* leads to edema formation, (k) *In situ* hybridization detected expression of *ift80* and *ift172* in limb buds of *X. laevis* (black arrowhead). po - posterior; a - anterior; pr - proximal; d - distal (l) Categorization of mutant froglets according to phenotypic strength of limb malformations.
Error bars indicate SEM. $p>0.05$ ns (not significant); $p<0.05$ *; $p<0.01$ **; $p<0.001$ ***
